# PKC Promotes T-Tubule Membrane Loss by Activating a PKD– NFκB Endocytic Pathway

**DOI:** 10.1101/2025.05.04.651469

**Authors:** A. Vilchez, A.-K. M. Pfeuffer, J. Weßolowski, D. J. Fiegle, P. Andrä, P. Potue, L. K. Küpfer, T. S. Shankar, C. H. Selzman, S. G. Drakos, C. Heim, T. Volk, T. Seidel

**Affiliations:** Institute of Cellular and Molecular Physiology, Friedrich-Alexander-University Erlangen-Nürnberg, Erlangen, Germany; Nora Eccles Harrison Cardiovascular Research and Training Institute, University of Utah, Salt Lake City, USA; Division of Cardiothoracic Surgery, University of Utah, Salt Lake City, USA; Division of Cardiovascular Medicine, University of Utah, Salt Lake City, USA; Department of Cardiac and Vascular Surgery, Klinikum Bayreuth, Medical Campus Oberfranken of Friedrich Alexander University, Bayreuth, Germany

## Abstract

**Background:** In heart disease, the membrane of the cardiomyocyte transverse-axial tubular system (TATS) deteriorates. This impairs contractility, hinders recovery and predisposes to arrhythmia. However, the key signals and cellular processes driving TATS loss are not understood. We investigated protein kinase C (PKC) and its downstream signals in animal and human cardiomyocytes.

**Methods:** Ventricular cardiomyocytes were isolated from healthy adult rat, rabbit and failing human hearts and treated with the PKC activator phorbol 12-myristat 13-acetat (PMA) or receptor-mediated agonists, alongside inhibitors targeting PKC, PKD, NFκB, MKK1–ERK1/2, NFAT, or endocytic pathways. TATS density was analyzed by confocal microscopy using lipophilic membrane dyes. Signaling pathway activation was determined by Western blotting and RNA sequencing. Ca^2+^ signals and contractility were assessed in rat cells. Mechanisms of TATS loss were studied using endocytosis assays involving fixable dextran.

**Results:** PMA induced severe TATS loss, which was prevented by inhibiting PKC, PKD, NFκB or MKK1, but not by blocking NFAT or p38 MAPK. Receptor-mediated PKC activation also decreased TATS density. All effective inhibitors suppressed IκB*α* expression. RNA sequencing indicated PMA-mediated activation of the NFκB and MAPK/ERK pathways and genes related to endocytosis. NFκB inhibition did not suppress the MAPK pathway, but MKK inhibition suppressed NFκB. PMA decreased Ca^2+^ transient amplitudes and contractility, whereas NFκB inhibitors preserved both. Dextran assays revealed that TATS membranes were internalized via a macropinocytic process that followed saturation kinetics was upregulated by PMA, downregulated by NFκB inhibition, and required PI3K, myosin I, and clathrin-independent endocytosis and correlated with the rate of TATS loss. Key findings were consistent in human cardiomyocytes and in ex-vivo rat and rabbit myocardial slice culture.

**Conclusions:** PKC activation drives TATS loss in human and animal myocytes via PKC–PKD–NFκB, T-tubules are degraded by endocytic internalization, offering a new perspective on how cardiomyocyte membranes may deteriorate in heart disease.

## 1. Introduction

Chronic heart failure (HF) affects millions of people worldwide and is associated with high hospitalization and mortality rates^1^. As a progressive clinical syndrome, it leads to adverse remodeling of the myocardium, which further impairs cardiac function, resulting in a vicious circle of maladaptive remodeling and disease progression. Despite advances in therapies, preventing or even reversing HF progression remains a major challenge and requires a better understanding of the cellular mechanisms and pathways that lead to pathological cardiac remodeling^2,3^.

One important function affected by the adverse cardiac remodeling in chronic HF, myocardial infarction and other heart diseases is excitation-contraction (EC) coupling^4^. EC coupling comprises the electrical excitation via an action potential, which elicits Ca^2+^ release into the cytosol from the extracellular space and intracellular stores. This triggers cardiomyocyte contraction and regulates its force and timing. Subsequently, cytosolic Ca^2+^ is removed to initiate relaxation. Ventricular cardiomyocytes possess numerous adaptations that enable them to constantly repeat this Ca^2+^ cycle very efficiently and effectively. An important structural adaptation is the transverse-axial tubular system (TATS or t-system), consisting of regularly arranged tubular invaginations of the cell membrane, most of which are aligned transverse to the longitudinal cell axis. The purpose of t-tubules is the creation of junctions between sarcolemmal L-type Ca^2+^ channels (LTCCs) and sarcoplasmic ryanodine receptors (RyR). This reduces the diffusion distance of Ca^2+^ from the extracellular space to RyRs, enabling rapid and synchronous Ca^2+^ release and contraction. Importantly, remodeling processes in HF reduce the density of t-tubules and change their shape and orientation^5–7^. This leads to an increased distance between RyR clusters and LTCCs, causing slow and inhomogeneous Ca^2+^ release. In this way, t-system remodeling contributes significantly to contractile dysfunction in HF ^8–10^. Furthermore, t-tubule loss has been associated with impaired cardiac recovery^7^ and arrhythmogenic events^11–13^.

Given these pathophysiological mechanisms, preventing or reversing t-tubule remodeling would be highly desirable. However, to date no suitable therapeutic strategies are available. To make this achievable, a much better understanding of the mechanisms and signaling pathways underlying t-system remodeling is needed. Although recent studies have found that certain proteins, triggers or signals are associated with t-tubule loss^14^, the specific details of t-tubule degradation, including signaling pathways and the actual cellular processes, remain elusive.

Regarding signaling pathways, several studies point towards inflammatory signals to be involved in t-system degradation. It has been reported that glucocorticoids may prevent t-tubule loss^15^. Glucocorticoids exert anti-inflammatory effects, e.g. by inhibiting NFκB signaling^16^. Another study found that protein kinase C (PKC) activation can accelerate t-tubule loss^17^. Interestingly, PKC is pro-inflammatory and activates NFκB^18^. In accordance with this, myocarditis has been linked to t-tubule alterations^19^. Moreover, PKC is generally considered to be activated in heart failure and cardiac ischemia. While positive effects of PKC have been reported, especially in the setting of ischemia-reperfusion injury^20,21^, chronic and over-activation of PKC have been linked mainly to negative effects on cardiomyocytes and the heart, especially in heart failure and cardiac hypertrophy^22–24^. This suggests a link between NFκB signaling and t-system loss.

Even less is known about the actual cellular processes that lead to remodeling and loss of the t-system. A number of triggering mechanisms have been suggested, such as cellular strain and mechanical overload^25–28^ or remodeling of the extracellular matrix, especially fibrosis^6,29–31^, as well as several proteins like junctophilin-2^32,33^, amphiphysin 2 (BIN1) ^34–37^, nexilin ^38^ or mitsugumin 53 ^5,39,40^. Also, autophagy, which is dysregulated in heart failure, has been linked to t-tubule loss^15^ as well as alterations of the cytoskeleton^17^. It is also known that detubulation is possible by application of sudden osmotic stress causing fast shrinkage and expansion of the cell^8^ or by disruption of phospholipid-protein interactions in the cell membrane^41^. Nevertheless, it is almost completely unknown how t-tubules spontaneously disappear in heart disease or in cell cultures.

The present study focuses on the effect of PKC activation on t-tubule loss in isolated rat and human cardiomyocytes, as well as in rat and rabbit myocardial tissue. We identify PKC and the downstream effectors protein kinase D (PKD) and NFκB as the major pathway that initiates the loss of t-tubules in cardiomyocytes. Furthermore, we identify activation of a novel macropinocytosis-like mechanism of t-tubule internalization as the underlying cellular process of t-system degradation.

## 2. Methods

### 2.1. Isolation and culture of rat and rabbit cardiomyocytes

The use of animals for all experiments in this study was approved by the local Animal Care and Use Committee of the University of Erlangen-Nürnberg. Cell isolation was carried out as previously described^15,42^. Female Wistar rats (150-250 g) were anaesthetized by intraperitoneal injection of 5-10 mg/kg xylazine and 80-100 mg/kg ketamine and then killed by cervical dislocation. Thoracotomy was carried out quickly to excise the heart, which was then connected to a Langendorff apparatus by aortic canulation. After 5 min of perfusion with a Tyrode-like solution containing 1 μM of free Ca^2+^, 160 U/mL collagenase CLS type II(Biochrom) and 0.6 U/mL protease type XIV(Sigma) were added and perfusion continued for 15 min. Finally, the heart was perfused for another 5 min without enzymes. Myocytes from the left-ventricular wall were then dislodged by mincing and gently agitating the digested myocardium. Myocytes were stepwise adapted to physiological Ca^2+^ levels and transferred to cell culture dishes containing culture medium M199 (Sigma, M4530), 0.1% BSA (Roth, 0163.2) and 1% Primocin (InvivoGen, ant-pm-1) or 100 mg/mL penicillin + 0.1 mg/mL streptomycin. During all procedures, the temperature was kept at 33-37° C. The cells were then distributed in petri dishes (Sarstedt, 83.3900), randomly assigned to the experimental groups, and kept in an incubator at 37 °C and 80% humidity and 5% CO_2_ until further use. Cardiomyocytes from rabbit hearts were isolated following the same protocol. New Zealand White rabbits, age 10-12 weeks (2.5-3.5 kg), were anesthetized by i.m. injection of 25 mg/kg ketamine and 5 mg/kg xylazine. Effectiveness of anesthesia and analgesia were verified by the absence of pain reactions. 200 mg/kg of pentobarbital sodium were then injected into the dorsal ear vein via a catheter, leading to deep anesthesia and cardiac arrest. The heart was removed quickly and transferred into a 4 °C cardioplegic solution^43^ containing 80 mM potassium glutamate, 10 mM NaCl, 30 mM 2,3-butanedione monoxime, 50 mM sucrose, 25 mM KH_2_PO_4_ 5 mM MgSO_4_, 1 mM CaCl_2_, 1 mM allopurinol, 5 mM adenosine, 5 mM glutathione, and pH 7.4 to the lab, where myocytes were isolated as described for rat hearts.

### 2.2. Isolation and culture of human cardiomyocytes

Left-ventricular human tissue samples were obtained from explanted hearts via the Cardiac Transplant Program of the University of Utah Health Science Center, the Department of Cardiac Surgery at Bad Oeynhausen, Germany, and the Cardiac Transplant Program at the University Hospital Erlangen, Germany. Additional samples from human hearts were also collected during left-ventricular assist device implantation at the University Hospital Erlangen, Germany. The corresponding institutional review boards approved the study, and all patients provided written informed consent, according to Declaration of Helsinki principles. The isolation of human cardiomyocytes was performed following a published protocol^44^. In brief, the tissue was cut with a vibratome into 300 µm thick slices and digested with 1 mg/mL proteinase XXIV (Sigma Aldrich, P5147) for 12 min, followed by 4 mg/mL collagenase CLS type I (Worthington LS004196) with 5 µM CaCl_2_ in a solution containing 20 mM KCl, 10 mM KH_2_PO_4_, 10 mM MgCl_2_, 10 mM glucose, 70 mM glutamic acid, 20 mM taurin, 10 mM beta-hydroxybutyrate, 30 mM 2,3-butanedione monoxime (BDM) and 2 mg/mL bovine serum albumin. The digested slices were then transferred into a solution containing in 20 mM KCl, 10 mM KH_2_PO_4_, 10 mMMgCl_2_, 10 mM glucose, 70 mM glutamic acid, 20 mM taurin, 10 mM beta-hydroxybutyrate, 5 µM CaCl_2_, 30 mM BDM and 10 mg/mL bovine serum albumin and dissociated with forceps. The Ca^2+^ concentration was slowly increased in ten steps to a physiological level. Undigested tissue fragments were removed by filtering the cell suspension through a nylon mesh with 180 μm pore size, and the solution was replaced by culture medium M199 (Sigma, M4530) supplemented 1mg/mL with BSA and 100 mg/mL penicillin / 0.1 mg/mL streptomycin. Pharmacological agents or the respective vehicles (ethanol, DMSO) were added, and the isolated cells then cultured for one day at 37°C and 5% CO_2_.

### 2.3. Preparation and culture of rabbit and rat myocardial slices

Female New Zealand White rabbits (2.5–3.5 kg) were sedated, euthanized and the heart removed in the same way as described for cell isolation. Myocardial slices from the left ventricle were prepared following published protocols^43,45^. Tissue slices were cultured for six days in tissue culture systems (MyoDish MD-1.1, InVitroSys) in culture medium, containing M199 (Sigma M4530), supplemented with 10 ng/mL insulin, 5.5 µg/mL transferrin, 6.7 ng/l selenium, 50µM β-mercaptoethanol, 100 U/mL penicillin + 0.1 mg/mL streptomycin and 20nM cortisol 20. All slices were slightly stretched to a diastolic preload of 1000 to 1500 µN and paced at 0.5 Hz. PMA at a concentration of 50 nM or ethanol as control were added after 24 h in culture. Two thirds of the medium were exchanged every 48 h.

Rat myocardial slices were prepared and cultured similarly. Immediately after excision, the heart was immersed in a cold storage buffer, containing 0.5 mM CaCl_2_, 30mM BDM, 10mM glucose, 10mM HEPES, 5.4 mM KCl, 2mM MgCl_2_, 138 mM NaCl, 0.33 mM NaH_2_PO_4_ and pH 7.4. After storage periods of 30 min to 2 h, the right ventricle as well as the base and apex of the left ventricle were removed and the remaining left ventricle, including the septum, placed in a petri dish with the basal site facing down. It was poured over with liquid low-melt agarose (Roth, 6351.5) at 37 °C. The chamber of the ventricle was also filled with agarose to improve stability during the subsequent cutting on a vibratome (Leica, VT1200). Ring-shaped slices of 300 µm thickness of the LV were created and cut in half to obtain slices from the LV wall or the septal wall with visible myocyte orientation. Length and width of the slices were 5-8 mm and 3-4 mm, respectively. The slices were then glued to plastic holders, mounted in culture chambers (InVitroSys). A preload of 500-750 µN and a pacing rate of 0.5 Hz were applied for 6 d. After 24 h, 10 nM PMA or ethanol as control were added and kept in the medium until the end of culture. Culture medium and other conditions were the same as for rabbit slices.

### 2.4. Isolation of rat cells from myocardial slices

To obtain cardiac myocytes from cultured rat cardiac slices electrical stimulation was discontinued and 30 mM BDM added to the medium. The slices were transferred into a petri dish, the triangular plastic anchors were removed and the slices were subjected to the same isolation protocol as described for the human cardiomyocyte isolation^44^.

### 2.5. Western Blots

After culture and treatment, cardiac myocytes were collected in their respective media and centrifuged at 2000 rpm for 2 min. The supernatant was discarded and the cell pellet stored at -80 °C until use. The frozen pellets were resuspended in 250 µL of TNE-buffer and homogenized by pipetting, then sonicated (3 *×* 5 min, on ice). The samples were incubated for 30 min on ice and then centrifuged for 10 min at 13.000 rpm at 4 °C. The resulting supernatant was collected and its protein concentration quantified using a BCA assay (Pierce kit, 23227, ThermoFisher). Electrophoresis was performed on 10% polyacrylamide gels on ice at 25 mV for 1.5 h, after which proteins were transferred to a PVDF membrane, using a Trans-Blot SD Semi-Dry Transfer Cell (BioRad, US) for 1 h. The transfer membrane was incubated in blocking solution at room temperature for 45 min before incubating with the primary antibody. After this, the membrane was washed (3 *×* 5 min) with TBS buffer and subsequently incubated with the secondary antibody. Signal was detected with a SuperSignal West Femto kit (34096, Thermo Fisher, Germany). A table specifying each antibody and the solutions is available in the Supplementary Methods.

Western blot results were quantified with Fiji/ImageJ software. After verifying that the image was not saturated, noise-filtering was applied by Gaussian Blur (sigma = 1) and background was subtracted with a rolling ball radius of 50 pixels. On the resulting images, signal intensity of the protein bands (*S*_i_) was measured and normalized to the total intensity of the Ponceau-stained bands of their respective lane (*P*_i_) and then to the reference band. Because the PMA-treated groups exhibited the highest signal in most experiments, PMA was used as the reference group to reduce noise artefacts^46^. Thus, the resulting normalized value for each band *i* was calculated as *S*_i_ / *P*_i_ / (*S_PMA_ / P*_PMA_).

### 2.6. RNAseq

After cell isolation and culture, the viability of cardiomyocytes was verified by light microscopy. If the fraction of rod-shaped myocytes with visible cross striations was at least 70%, cells were used for RNA sequencing, otherwise they were excluded.

Freshly isolated cardiomyocytes were subjected to their respective treatments and cultured; inhibitors were incubated for 1 h prior to PMA addition and then 4 h together with PMA for a total of 5 h, all vectors were accordingly added to the respective controls. Cells were collected in their respective media and centrifuged at 2000rpm for 2 min, after which the supernatant was discarded and the pellet was frozen and stored at -80 °C until use.

Total RNA was extracted using a commercial kit (Nucleospin, Machery Nagel, Germany) according to the manufacturer’s instruction. Samples from one isolation were handled in parallel to minimize batch-effects between groups. After extraction purity was controlled by measuring 260/280 and 260/230 absorbance ratios by spectrophotometry (DS-11+ Spectophotometer, DeNovix, Wilmington USA). All samples contained more than 100ng of RNA. RNA was then frozen at -80°C until handing them over to the sequencing facility. A second quality check using a commercial device (BioAnalyzer, Agilent, Santa Clara, USA) was used to exclude samples with degraded RNA.

The RNA was processed into a library with the Illumina Stranded mRNA Kit (Illumina, San Diego, USA). The library was sequenced as 150+150bp paired-end reads on an Illumina Novaseq 6000 platform. Each sample yielded 32-57 million raw reads. After removal of optical and proximal duplicates, adapter and quality trimming, the resulting reads were aligned against the mRatBN7.2 reference genome with the STAR aligner software (version 2.7.10 a)^47^. Transcript quantification was performed with Salmon 1.9 software. Statistical analysis of normalized pseudocounts was carried out with the DEseq2 package (v1.44) in the R statistics environment (v4.4.2) library.

#### 2.6.1. Pathway Analysis

The PROGENy package (v1.26) was employed for measuring pathway activity^48^. Our samples were compared to the top 2000 genes of the rat and human model for each pathway. Log-transformed normalized counts and the Wald statistic, respectively, were used for analysing activity in single samples and in grouped comparisons. Prior to the analysis, our data was processed to filter out any invalid (nan) or duplicate entries, so as not to introduce errors into the statistic. The results were displayed in heatmaps and scatterplots created using the pheatmap (v1.0.12) and ggplot2 (v3.5.1) packages, respectively.

#### 2.6.2. GSEA

Gene set enrichment analysis (GSEA) was performed using the list of differentially expressed genes from the DESeq2 analysis. Invalid entries (nan) and duplicates were removed. Genes were then ranked in descending order by their stat values (log_2_-fold change divided by its standard error). An fgsea (v1.30) was performed using a number of preselected gene sets obtained from the Molecular Signatures Database (MSigDB v2024.1.Hs/Mm)^49,50^ and 10^6^ permutations. Gene sets were selected based on a keyword search in curated (C2) gene sets. The results fitting best to keywords and pathways (NFAT, Ca^2+^ signaling, cardiac muscle contraction, cardiomyopathy, endocytosis, integrins, cytoskeleton) were used. Where available, mouse gene sets were used, otherwise human. A gene set with t-tubule associated genes was created by merging the lists from literature research and immunostaining studies, curated by Cheah et al.^51^ The normalized enrichment score and adjusted p-values were plotted using the ggplot2 package (v3.5.1).

#### 2.6.3. Mechanism-related genes

Genes were selected if their expression level was significantly altered (adjusted p-value < 0.05), upregulated when comparing PMA with CTRL (log_2_-fold change > 0) and downregulated when comparing PMA+TN with PMA (log_2_-fold change < 0), or vice versa, i.e., log_2_-fold change (PMA vs. CTRL) < 0 and log_2_-fold change (PMA+TN vs. PMA) > 0. To obtain lists of genes associated with different processes, various gene databases were used. The KEGG (Kyoto Encyclopedia of Genes and Genomes) database was employed to identify genes associated with “membrane trafficking” and “cytoskeleton proteins”; specifically, the BRITE pathway “BR: hsa04131” was used for membrane trafficking, and “BR: hsa04812” for cytoskeleton proteins. For genes related to endocytosis, the keyword “endocytosis” was used in the Gene Ontology (GO) database, filtered for Rattus norvegicus, which yielded 737 genes. The differentially expressed genes were then compared to the genes retrieved from the database search and visualized. Data processing was performed using R and RStudio (Version 2021.09.0).

### 2.7. Staining protocols

#### 2.7.1. Membrane Staining of living cells

Membranes of living cells were stained for t-system analysis with 8.5 µM of the lipophilic dye Di-8-ANEPPS (Sigma-Aldrich, D1065) or, in specific occasions, 16.5 µM of the lipophilic dye FM4-64 (Thermo Fisher, T13320). The nucleus was stained with Hoechst 33342 (Invitrogen, H3570). Cells were transferred to a 1.5 mL tube while still in their culture media and, together with the dyes, 30mM of 2,3-butanedione monoxime (Thermo Fisher, A14339) was added to avoid contraction during image acquisition. Imaging was always completed within 4 h after staining.

#### 2.7.2. Dextran assay

After culture, living cardiomyocytes were transferred from the cell culture dish to a 1.5 mL tube and mildly centrifuged (80 *× g* , 30 s). Excess culture medium was removed, leaving a residue of 50 μl. 25 mg/mL of fixable dextran conjugated to Alexa Fluor 488 (10 kDa, Thermo Fisher, D22910) were incubated with the lid open for 5 – 80 min at 37 °C, 80% humidity and 5% CO_2_. If not declared differently, the incubation time was kept at 10 min. In experiments containing a wash-out of dextran, cells were then placed back into normal culture medium and incubated for 10 – 40 min at 37 °C, 80% humidity, and 5% CO_2_. Then, a washing step with a phosphate-buffered solution containing 130 mM NaCl, 5.4 mM KCl, 0.5 mM MgCl_2_, 0.4 mM NaH_2_PO_4_, 22 mM Glucose, 25 mM HEPES, 5.8 mM NaHCO_3_ and 2 mM CaCl_2_ at 4 °C followed. Afterwards, the fixable lipophilic dye AM4-66 (Biotium, 70050) was added at a concentration of 1 µM for 5 min on ice. Subsequently, after gentle centrifugation (80 *× g*, 30 s), the supernatant was discarded, and the myocytes were fixed with 2% PFA for another 5 min. In the final step, the supernatant was removed after further centrifugation, and the cells were stored in PBS on ice and promptly imaged on a confocal microscope.

#### 2.7.3. Immunofluorescence of isolated cells

Immunostaining was carried out in 1.5 – 2 mL tubes. Briefly, 1.5 mL of suspended cells were transferred to the tube and gently centrifuged for 45 s at 80×g, after which the supernatant was carefully aspirated and discarded, and the fixative was added. Following pertinent timing, the fixing agent was removed and the cells incubated with PBS for 10 min, and after removal of PBS the cells were incubated with the primary antibodies for 4-5 h at room temperature or overnight at 4 °C. The samples were then washed with PBS (3 *×* 10 min) before adding the secondary antibodies, which were incubated for 3 h at room temperature. Finally, the cells were washed with PBS for 3 *×* 10 min and stored at 4 °C until imaging. A table with the materials and details of each staining is provided in the Supplementary Methods.

#### 2.7.4. Immunofluorescence of tissue slices

As described recently^45^, rabbit tissue slices were fixed in 2% PFA for 10 min immediately after culture and then washed 3 *×* 5 min in PBS. Primary antibodies against RyR2 (IgG1, mouse, C3-33, Thermo Fisher) diluted 1:200 in blocking solution (5% normal goat serum, 5% BSA, 0.25% Triton-X in PBS) were added and incubated for at least 4 h at RT or overnight at 4 °C. After washing with PBS, the secondary antibody goat anti-mouse IgG1 AF-488 (Thermo Fisher A-21121) was added in blocking solution and incubated for at least 3 h RT. After washing, slices were incubated with 40µg/mL wheat germ agglutinin (WGA-AF-647 Thermo Fisher W32466) and 1.67µg/mL DAPI (3665, Roth, Karlsruhe, Germany) in PBS for another 3 h. Slices were embedded in Fluoromount G (00-4958-02, Thermo Fisher) on a microscope slide, covered with a coverslip and let dry for 1–7 days at 40–45% humidity before imaging.

### 2.8. Confocal microscopy

#### 2.8.1. Isolated cells

Three-dimensional confocal imaging of living cells stained with Di8-ANEPPS and Hoechst was done as described before^15^. Briefly, the cell suspension was imaged with an LSM780 confocal microscope (Zeiss), using a 63*×* oil immersion objective and a voxel size of 0.1*×*0.1*×*0.2 µm^3^. Each stack consisted of at least 1280*×*384*×*50 voxels, yielding an image with a volume of 128*×*38.4*×*10 µm^3^ per cell. Human dextran-stained cells were acquired with a Leica SP8 confocal microscope, an image size of 1536*×*384*×*50 voxels, and an additional acquisition channel was introduced (wavelength from 580 to 620 nm) to detect lipofuscin granules^45^. In human cells stained with Di8-ANEPPS lipofuscin granules were excited by a 633 nm laser and were detected with an additional channel (700-800 nm). In the case of fixed immunostained cells, the stack height was reduced to 5µm and a region in the center of the cell was imaged. Except for few pilot experiments, the researchers were blinded against the sample groups during image acquisition. Cardiomyocytes and tissue regions were selected with light microscopy to avoid any bias regarding the t-system density. Attenuation correction was applied by adjusting the laser power with depth and subsequent image-intensity correction^52^. A table specifying imaging parameters for each fluorophore can be found in the Supplementary Methods.

#### 2.8.2. Tissue Slices

Rabbit cardiac tissue slices were imaged as described earlier^43,45^. A one-track protocol was used, with parallel excitation of DAPI, AF488 and AF-647 by laser wavelengths of 405, 488 nm and 633 nm, respectively. The size of each image stack was 1496*×*1496*×*120 voxels with of 0.1*×*0.1*×*0.2 µm^3^ voxel size. Depth-dependent laser power increase was applied to compensate for signal attenuation. Imaged tissue regions were chosen randomly by light microscopy by a researcher blinded against the sample groups.

### 2.9. Ca^2+^ imaging and analysis

#### 2.9.1. Epifluorescent Imaging of cardiomyocytes with Fura-2

Rat cardiomyocytes were gently centrifuged at 80 *× g* for 3 min and the culture medium was then removed. Cells were incubated in an extracellular solution containing 138 mM NaCl, 4 mM KCl, 1 mM MgCl_2_, 0.33 mM NaH_2_PO_4_, 10 mM glucose, 10 mM HEPES, 2 mM CaCl_2_ and pH 7.40 (adjusted with NaOH) containing 1 µmol/L Fura-2-AM and 0.1% pluronic acid at room temperature for 20 min. Subsequently, cells were washed and resuspended in 1 mL of extracellular solution and incubated for an additional 10 min. Afterwards, imaging was performed in a small dish with a glass bottom on an inverted microscope (Leica, Germany), with continuous superfusion of extracellular solution (160-250ml/h) at 34-35 °C. Field stimulation was applied using a voltage generator (MyoPacer, IonOptix, USA). Cells were excited at 340 and 380 nm with a switching rate of 200 Hz (Hyper-switch, IonOptix, USA). The fluorescence intensity ratio at 340/380 excitation was used as measure of the free intracellular Ca^2+^ concentration. Cells were pre-paced for at least 1 min at 1 Hz before recordings. The sarcomere length was assessed by the frequency transform of the transmitted light image using IonWizard 6.6 software package (IonOptix, Westwood, MA, USA). Electrical stimuli and temperature were recorded. Immediately after pre-pacing, the cells were paced with increasing frequencies of 0.5, 1, 1.5, 2, 2.5, 3, and 4 Hz. Each interval consisted of at least 24 stimulations and at least 12 sec.

The resulting time-series data was analysed with custom-written Matlab scripts (Mathworks, USA, version 2022a and higher). First, peak detection of recorded voltages yielded stimulation times. The raw signals (F340/380 ratio and sarcomere length) were filtered with a moving median filter (radius 10) and a moving mean filter (radius 5). The minimum value of each time series (offset) was subtracted and the noise amplitude (A_noise_) was detected by searching the maximum signal within the 200 ms interval preceding a stimulus at pacing frequencies ≤ 1 Hz. Next, to identify individual Ca^2+^ signals or contractions, peak detection was applied to the corresponding signals, using a minimum peak height of 2 • A_noise_. If no peak was detected, the stimulus was identified as not captured. Otherwise, the signal was further analysed to quantify its diastolic level, amplitude, time to peak, time to baseline and Ca^2+^ transient duration. Parameters from 6 successive stimulations at the end of an interval of a certain stimulation frequency were averaged and used as representative value for one cell at the respective frequency.

#### 2.9.2. Confocal line scans of cardiomyocytes including 3D t-system quantification

Confocal line scans were carried out as described earlier in detail^15^. Rat cardiomyocytes were stained with Hoechst for the nuclei, 8.5 µM Di8-ANEPPS for the membranes and loaded with 10 μM Fluo-4 AM (Thermo Fisher, F14217) in extracellular solution containing 130 mM NaCl, 5.4 mM KCl, 0.5 mM MgCl2, 0.4 mM NaH2PO4, 22 mM glucose, 25 mM HEPES, 2 mM CaCl_2_ and pH 7.4 (adjusted with NaOH) and incubated for 20 min at room temperature. Cells responding to the electrical point stimulation were selected randomly by light microscopy by a researcher blinded against the sample group and paced by a standardized protocol via field stimulation. Image acquisition was performed in a bath containing extracellular solution with a Zeiss LSM 780 inverted confocal microscope, using a 63*×* oil immersion lens (NA 1.4), a 488 nm argon laser for excitation, and emission filters set to 489-550 nm (Fluo-4) and 566-718 nm (Di8-ANEPPS). Pixel size was 0.1*×*0.1 μm^2^. One line contained 1024 pixels. After adjusting the focal plane to the cell centre, the cell was stimulated for 30 s at 1 Hz prior to image acquisition. After a pause of 25 s, the myocyte was stimulated and a line scan simultaneously recorded in the cell centre along the myocyte long axis (1024 pixels, pixel dwell time 1.26 μs, sampling rate 529 Hz = 1.89 ms per line). During the 25s pause, a 2D scan (1024*×*384 pixels, pixel size 0.1*×*0.1 µm^2^) and a 3D scan (1024*×*384*×*23 voxels, voxel size 0.1*×*0.1*×*0.3 µm^3^) was carried out to obtain 2D and 3D information on the t-system density around the subsequently scanned line. For each data set, 2000 lines were scanned, corresponding to 3.78 s.

Line scans were noise-filtered, corrected for spillover from the Di8-ANEPPS channel into the Fluo-4 channel and analysed using a custom-written Matlab script (R2022b and higher, Mathworks). The nucleus was excluded from the analysis. For each pixel, a peak detection was carried out in its corresponding time series and a sigmoidal curve fitted, using parameter bounds derived from the peak detection and the sum signal of the line. In the fitted curve the time of maximum upstroke velocity was identified as activation time (t_a_). Using the membrane signal from the 3D stack, the 3D Euclidean distance of each pixel in the line to its closed t-tubule was calculated.

### 2.10. Image processing

ConfocaI image stacks were noise-filtered and deconvolved with measured point spread functions^53,54^. Subsequent image processing and analysis was performed with custom software as previously published^15,31,45,52,55,56^. In brief, all microscopy channels were segmented by applying histogram-based local thresholds by calculating the local standard deviation *s* and image mode *m* and the threshold *t* determined as t = c *• s* + m. The resulting binary images were created by identifying voxels with values ≥ t as signal, otherwise as background. A median filter with radius 1 was then applied to remove speckles. The 3D Euclidean distance transform (distance map) of the membrane and t-tubules served as measure of t-system density. High TT (t-tubule) distance indicated low t-system density, while low TT distance indicated high t-system density. For the single cells, the segmented signal of the membrane dyes Di8-ANEPPS, AM4-66, FM4-64 were used. In tissue slices, WGA was used. Cell masks were created by watershed-based segmentation and the use of a myocyte marker (ryanodine receptors) in tissue and manual segment merging in cells. This allowed to restrict all analyzes to the cytosol (Fig. 1A).

**Fig. 1:**
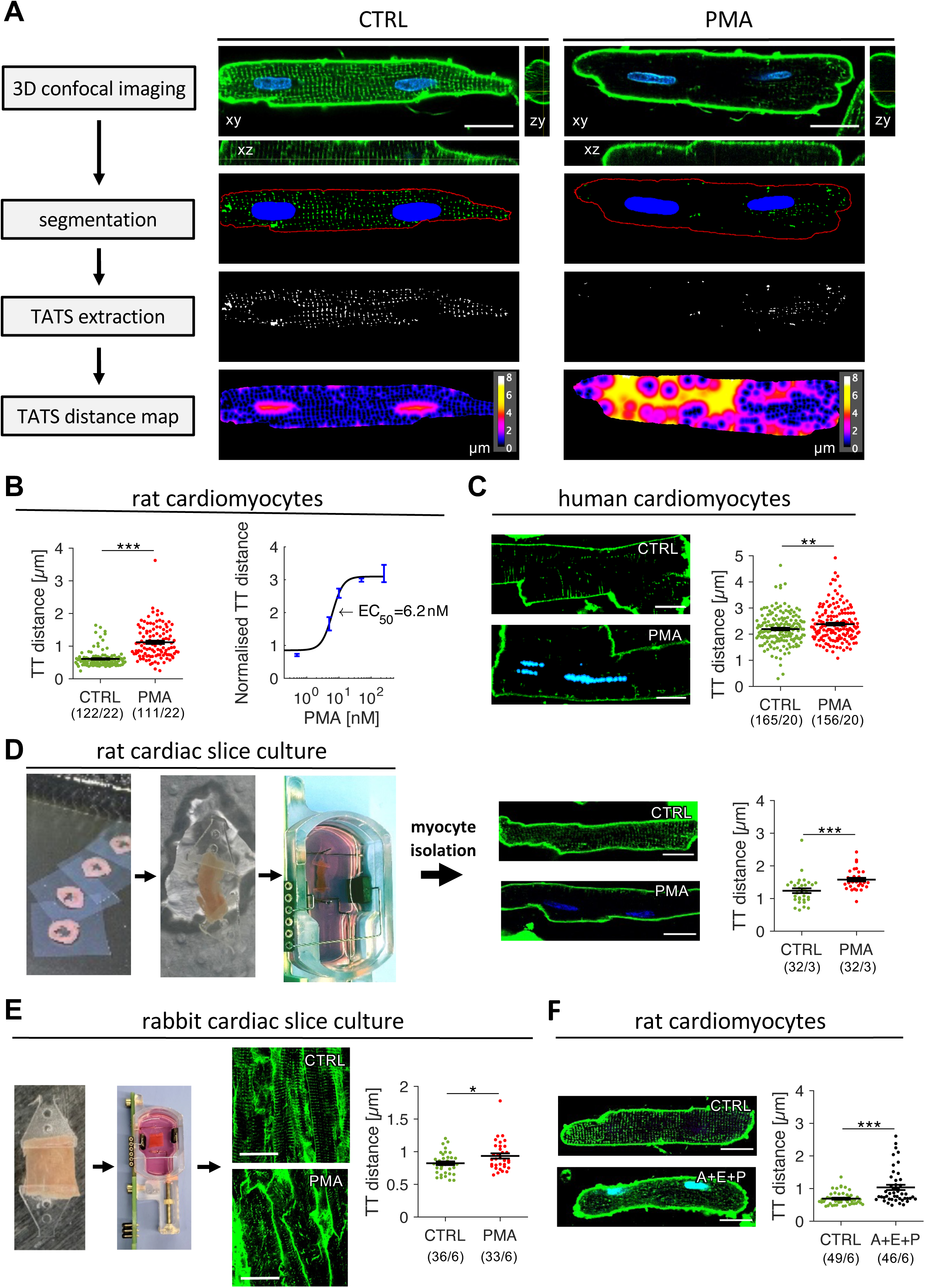
PMA induces TATS loss in cardiomyocytes. **(A)** Three-dimensional images of cells with Di-8-ANEPPS membrane staining and Hoechst nucleus staining were acquired with a confocal microscope (top) and then processed to segment the nuclei (blue), TATS (green) and surface membrane (red). Subsequently, the TATS was extracted and the Euclidean distance map created to calculate the mean distance from each point inside the cell to the nearest t-tubule (TT distance). A control (CTRL) and a PMA-treated rat cardiomyocyte are shown. **(B)** Mean TT distance of isolated rat cardiomyocytes cultured for 24 h under CTRL conditions (n/N = 122/22) or with 50 nM PMA (111/22). The half maximal effective concentration (EC50) was determined by normalising the mean TT distance of PMA-treated cells to the CTRL value. PMA concentrations were 0.5 nM (n=16), 5 nM (n=16), 10 nM (n=32), 50 nM (n=234), and 250 nM (n=8). **(C)** Mean TT distance of left-ventricular cardiomyocytes isolated from failing human hearts, cultured for 24 h under CTRL conditions (165/20) or with 50 nM PMA (156/20). Example images of each group are shown, with Di8-ANEPPS staining (green) and detected lipofuscin highlighted (cyan). **(D)** Left-ventricular tissue sections from rat were kept in biomimetic culture and treated with vehicle (CTRL) or 10 nM PMA for 4 days. Living myocytes were then isolated and stained with Di-8-ANEPPS to visualize the TATS and analyse mean TT distance (32/3 in each group). **(E)** Left-ventricular slices from rabbit were kept in biomimetic culture and treated with vehicle (CTRL) or 50 nM PMA for 5 days. After culture, tissue slices were fixed, stained with WGA and imaged by 3D confocal microscopy to visualize the TATS and quantify mean TT distance (image volumes/animals: CTRL 36/6, PMA 36/6). **(F)** Mean TT distance of isolated rat cardiomyocytes cultured for 48 h under CTRL conditions (49/6) or with 1µM angiotensin II, 100 nM endothelin 1 and 10 µM phenylephrine (A+E+P, 46/6). All scale bars: 20 µm, * p< 0.05, ** p< 0.01, *** p<0.001, Unpaired non-equal variance t-test. Numbers below graphs indicate the number of cells or data points / the number of cell isolations (hearts).

Clusters of dextran, JPH2, RyR2 and BIN1 were detected by connected component analysis. The volume ratio of these signals, including dextran, was calculated by dividing the volume of dextran by the volume of the myocyte. Centroids of the connected components were used to calculate the centroid density (number of clusters were unit volume).

In most cells, the nucleus (stained either by Hoechst or by DAPI) was excluded from the analysis. In human cardiomyocytes we additionally excluded lipofuscin granules. Lipofuscin was detected by an additional channel, in which the fluorophores were only weakly present (see Fig. 1C), utilizing the broad emission spectrum of lipofuscin^57,58^.

### 2.11. Statistics

The data in the graphs are presented as mean ± standard error unless otherwise indicated. Where applicable (e.g. multiple measurements within the same slice or cell) a paired t-test was used. Otherwise, an unpaired two-tailed t-test with unequal variance assumption (Welch’s t-test) was applied, as indicated in the figure legends. The number of data points (n) and biological replicated (N) are provided in the respective graphs below the group names. When showing multiple comparisons within one chart or figure, the multiple comparison correction of p values according to Holm-Bonferroni was applied. The level of significance was set to α = 0.05.

Linear regression was performed, using the ‘fitlm’ function of MATLAB (Mathworks, version 2022b). The model was statistically assessed using an F-test versus a degenerate model with only a constant term.

RNA sequencing data was analyzed using the Wald test, and multiple testing correction was performed using the Benjamini-Hochberg method with a false discovery rate (FDR) threshold of 5%.

## 3. Results

### 3.1. PKC activators induce t-tubule loss across different models

To investigate the effects of chronic PKC activation on the t-system, we cultured isolated rat cardiomyocytes for 24 h in the presence of 50 nM PMA or vehicle as control (CTRL). Living cardiomyocytes were then stained with Di8-ANEPPS and imaged by confocal microscopy to acquire 3D images of the cell membrane including t-tubules. Hoechst DNA stain was used to visualize the nuclei. Fig. 1A shows an example from each group. We found that PMA strikingly reduced the density of t-tubules, with some cells displaying a nearly complete loss. To quantify t-tubule density, the membrane signal was converted into a binary image by threshold application. The t-system signal was extracted by removing the surface membrane. Subsequently, a distance map was created to obtain the 3D distance from each intracellular voxel to the closest t-tubule (distance map in Fig. 1A). It is well visible that the PMA-treated example cell contains regions of much higher distance to the closest t-tubule than the CTRL cell. Therefore, the mean t-tubular (TT) distance was used as a measure of t-tubule loss.

Quantification of TT distance in 111 PMA and 122 CTRL cells showed a nearly 2-fold increase in PMA cells (0.61±0.03 vs 1.14±0.07, p<0.01), indicating severe t-tubule loss (Fig. 1B). By investigating the effects of PMA on TT distance at different concentrations we obtained the dose-response relationship, which yielded a half maximal effective concentration (EC50) of 6.2 nM. This shows that 50 nM PMA elicited the maximum effect. Thus, this concentration was used in all subsequent experiments unless otherwise stated.

To determine if t-tubule loss can also be induced in human cardiomyocytes by PMA, we isolated primary myocytes from left-ventricular tissue samples obtained from human failing hearts and followed the same approach. Compared to rat cells, human cardiomyocytes presented much higher TT distance already at the control level, most likely a result of t-tubule loss resulting from heart failure^7^, but also of species differences^59^. Nevertheless, PMA still caused an additional loss of t-tubules after 24 h of incubation, although the difference between control and PMA-treated cells was lower than in rat (TT distance in µm, 2.22±0.08 in CTRL vs 2.42±0.1 in PMA, p<0.01, Fig. 1C). To explore the effect of PMA on the t-system in a third species, we used rabbit isolated cells. In consistency with rat and human myocytes, TT distance increased after PMA treatment (Fig. S1).

To verify whether t-tubule loss could be replicated in living myocardium in vitro, we cultured rat left-ventricular slices from rat hearts, using a biomimetic culture setup (Fig. 1D, left), in which slices were continuously paced to generate regular contractions, resembling physiological conditions. After one day in culture, tissue slices were treated with 10 nM PMA or EtOH (vehicle) as control. After a treatment period of 4 d, single cells were isolated from the slices and stained with Di-8-ANEPPS to assess t-system integrity as done after cell culture. Cardiomyocytes from slices treated with PMA showed a marked t-tubule loss (TT distance 1.58±0.07 µm vs 1.24±0.05 µm in CTRL, p<0.001, Fig. 1D, right).

To evaluate effects of PMA on the t-system without prior cell isolation, we repeated these experiments in rabbit myocardial slices, as t-tubule structure can be reliably visualized within fixed tissue in this species (Fig. 1E). Although the effect size was smaller than in rat cells after isolation from slices, significant t-tubule loss could be observed also in this model (TT distance 0.93±0.04 µm in PMA vs 0.84±0.03 µm in CTRL vs, p<0.05).

PMA causes acute and intense PKC activation by mimicking the effects of DAG, bypassing the receptor-mediated signaling cascade^60^. To investigate whether t-tubule loss can also be induced by receptor-mediated PKC activation, rat cardiac myocytes were treated for 24 h with a combination of substances acting via Gq-coupled receptors: angiotensin II, endothelin 1 and phenylephrine (A+E+P). A moderate yet significant t-tubule loss was observed in cells treated with A+E+P (TT distance 1.04±0.08 µm vs 0.69±0.03 µm in CTRL, p<0.005, Fig. 1F). We confirmed PKC activation in the A+E+P group by Western blotting of the PKC-specific phosphorylation of PKD at pS744-748 (Fig. S1C+D).

Collectively, these results show that PMA induces t-tubule loss in rat, rabbit and human cardiac myocytes and that these effects can be qualitatively reproduced in a multicellular ex-vivo model and by substances causing receptor-mediated PKC activation. The velocity of t-tubule loss may differ across species and appears faster in isolated cells than in intact myocardium.

### 3.2. PMA-induced t-system remodeling is mediated by PKC

PMA is considered a strong and specific activator of all classical (conventional) and novel PKC isoforms and also of the closely related protein kinase D (PKD)^61^. To verify the involvement of PKC in t-tubule loss and to shine light on the role of different PKC isoforms, we used broad-spectrum and isoform-specific inhibitors. First, to confirm that the effects of PMA on the t-system were a consequence of PKC activation, we used the broad-spectrum PKC inhibitors bisindolylmaleimide XI (BXI, 10 µM), a combination of 100 nM staurosporine and 500 nM sotrastaurin (S+S), GÖ6983 (GÖ, 1 µM), and bisindolylmaleimide 1 (B1, 1 µM), which inhibit both classical and novel PKC isoforms^62–65^. The addition of BXI or GÖ prevented t-tubule loss almost completely in rat cardiomyocytes (Fig. 2A). The effects of S+S and B1 were very similar, although slightly less pronounced. Western blotting with antibodies against the PKC-specific phosphorylation sites of PKD (pS744-748)^66,67^ confirmed the activation of PKC via PMA and the inhibition of PKC via the inhibitors (Fig. 2B). This suggests PKC activation as the main mechanism by which PMA induces t-tubule loss.

**Fig. 2:**
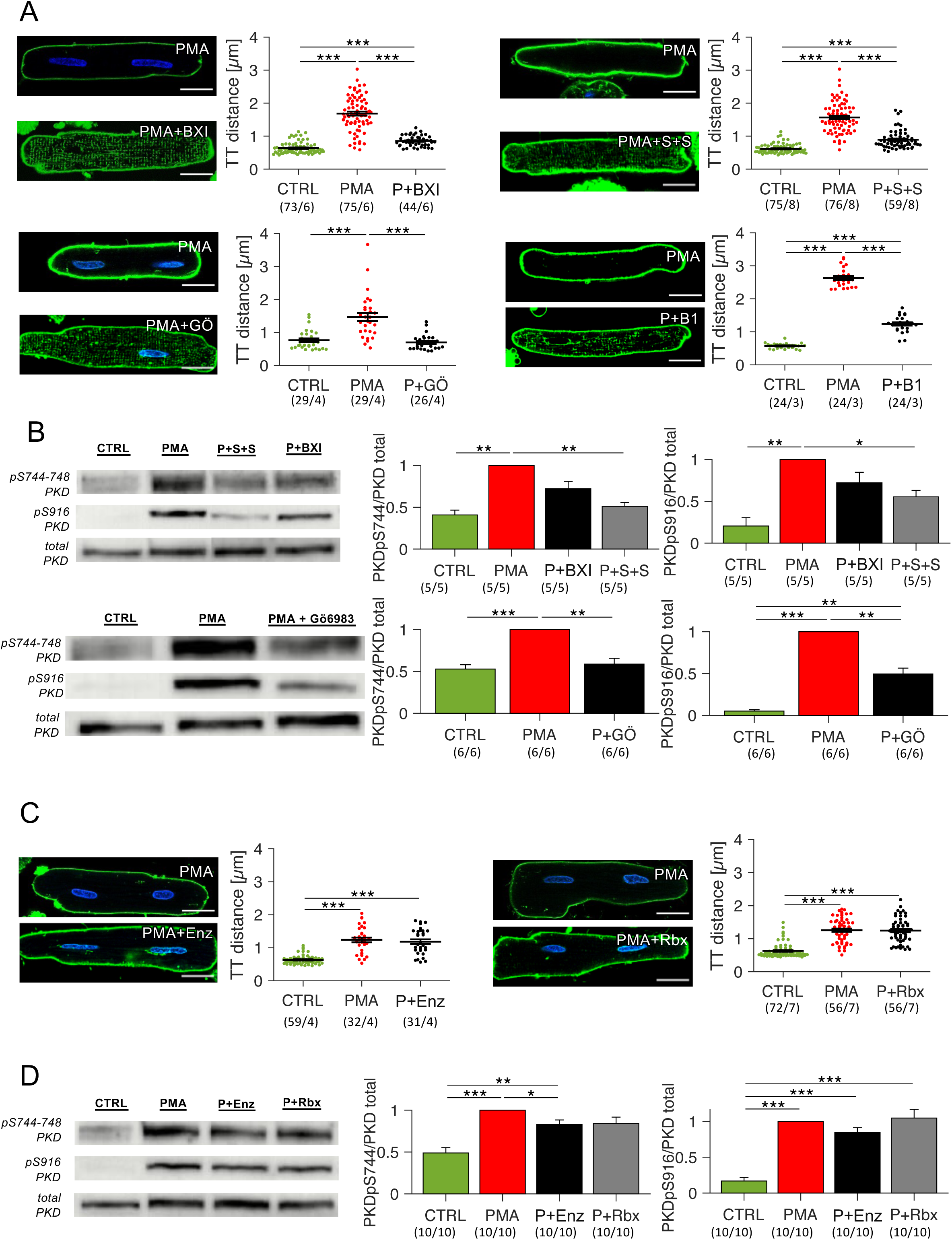
PMA induces t-system loss through PKC. **(A)** Effect of broad-spectrum PKC inhibitors on PMA-induced TATS loss. Cells were cultured for 1 h with the inhibitors bisindolylmaleimide XI 1µM (BXI), GÖ6983 1µM (GÖ), sotrastaurin 500nM + staurosporine 100nM (S+S), bisindolylmaleimide 1 500nM (B1). Then PMA was added. Vehicle (CTRL) and PMA alone served as controls. Example images of PMA- and inhibitor-treated cells stained with Di-8-ANEPPS are shown. Mean t-tubule distance was quantified and compared. In the case of BIS1, this was done at day 2 instead of day 1. **(B)** Effect of S+S, BXI and GÖ on PKD phosphorylation by PMA at S744-748 and S916, assessed by Western blotting. Band intensities were normalized to the total PKD intensity and then to CTRL on each blot. **(C)** Effect of PKCß inhibitors enzastaurin 300nM (Enz) and ruboxistaurin 50nM (Rbx) on PMA-induced TATS loss. Same experimental protocol as in (A). **(D)** Effect of Enz and Rbx on PKD phosphorylation by PMA at S744-748 and S916, assessed by Western blotting. Band intensities were normalized to the total PKD intensity and then to CTRL on each blot. Scale bars: 20 µm, * p < 0.05, ** p< 0.01, *** p< 0.001, unpaired, non-equal variance t-test, cells from matched cell isolations. Numbers below graphs indicate the number of cells or data points / the number of cell isolations (hearts).

We also investigated PKD phosphorylation at site S916, which indicates PKD activation by autophosphorylation (Fig. 2B). The results show that PKC activation and inhibition led to PKD activation and inhibition, respectively.

Because PKC*β* activation has been proposed as a pathomechanism in hypertrophy and HF^68,69^, we next applied two inhibitors with high specificity for PKC*β*. However, neither enzastaurin (300 nM) nor ruboxistaurin (50 nM) reduced t-tubule loss caused by PMA (Fig. 2C), even though it is likely that at the used concentrations they also inhibited other classical PKC isoforms^70^. Interestingly, enza- and ruboxistaurin had no significant effects on the phosphorylation of PKD at S744-748 and S916 (Fig. 2D), which would fit to PKD being a downstream target mainly of novel PKC isoforms^71^. These findings show that PMA-induced t-tubule loss is mediated via PKC and suggests a major role of PKD.

### 3.3. PKD activation downstream of PKC is essential for t-tubule loss

PKC is involved in a complex system of signaling processes and pathways (Fig. 3A). The best characterized downstream pathways are NFAT (classical PKCs), ERK (classic and novel PKCs) and NFκB (novel PKCs and PKD). To further elucidate the involvement of PKC/PKD and their downstream effectors in t-tubule loss, we applied multiple pathway inhibitors to rat cardiomyocytes in the presence of PMA.

**Fig. 3:**
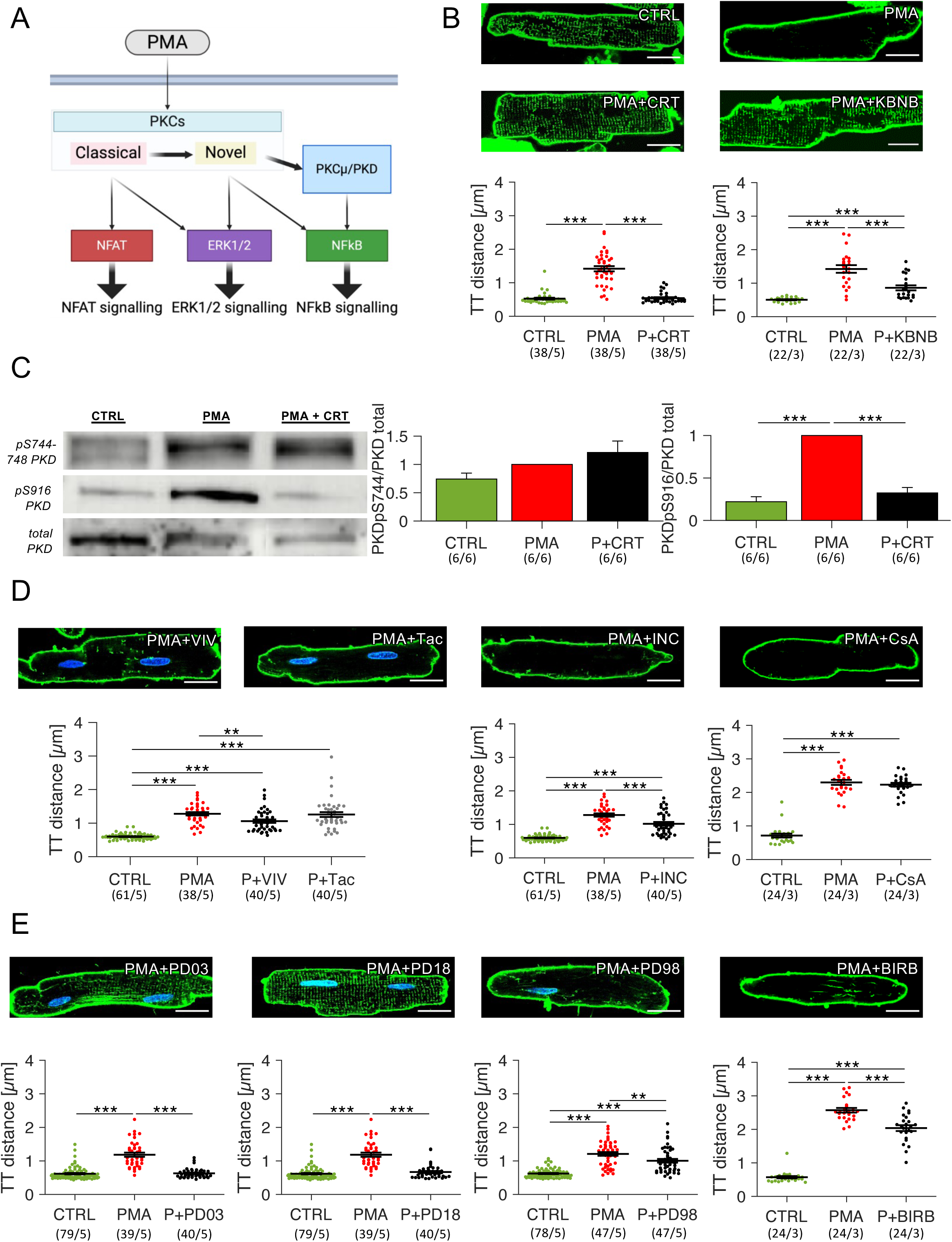
Effects of PKD, NFAT, and ERK inhibition on PMA-induced t-system loss. **(A)** Schematic representation of prominent downstream pathways of PKC; PMA activates PKC, whose isoforms contain classical (conventional), novel and atypical isoforms (atypical not shown). Downstream effectors and pathways are PKD (=PKCµ), NFAT, ERK1/2 and NFkB. **(B)** Effect of the specific PKD inhibitors CRT0066101 2.5µM (CRT) and KB-NB142-70 5µM (KBNB) on PMA-induced TATS loss, shown as t-tubular (TT) distance. Cells were cultured for 1 h with the inhibitors before addition of PMA and then cultured for another 24 h. Images show representative control (CTRL), PMA and PMA+inhibitor cells. **(C)** Phosphorylation of PKD at the PKC-specific (pS744-748) and autophosphorylation (pS916) site, assessed by Western blotting. Band intensities were normalized to total PKD levels. **(D)** Effect of four different NFAT inhibitors on PMA-induced TATS loss; cells were cultured for 24 h in the presence of PMA +/- tacrolimus 5µM (Tac), VIVIT 10µM (VIV), INCA 5µM (INC) or ciclosporin A 1µM (CsA). Cells treated with CsA were cultured for 48 h. Control cells (CTRL) were treated with vehicle. **(E)** Effect of four different ERK/MAPK pathway inhibitors on PMA-induced TATS loss; cells were cultured for 24 h in the presence of PMA +/- PD0325901 (PD03), PD184352 (PD18), PD98059 (PD98) or 48 h +/- BIRB796 (BIRB). Scale bars: 20 µm, * p < 0.05, ** p< 0.01, *** p< 0.001, unpaired, non-equal variance t-test, cells from matched cell isolations

First, we tested inhibitors that specifically target PKD, but not PKC. We used CRT0066101^72^ (2.5 µM) and KB-NB142-70^73^ (5 µM). Both inhibitors prevented PMA-induced t-tubule loss. Keeping t-system density at control level, this effect was most pronounced with CRT (*Δ*TT in µm, 0.53±0.03 in CTRL, 1.43±0.18 in PMA vs 0.54±0.02 in PMA+CRT, 0.87±0.08 in PMA+KBNB, Fig. 3B). The specificity of CRT was confirmed by Western blots showing that the phosphorylation at the PKC site s744-748 of PKD was unaffected, whereas its autophosphorylation site s916 showed significantly less phosphorylation (Fig. 3C) in the presence of PMA+CRT. This suggests that the effect of PKC on the t-system is predominantly mediated by downstream activation of PKD. To further test this hypothesis, we applied several inhibitors known to interfere with NFAT signaling: tacrolimus (5 µM), INCA-6 (5 µM) and VIVIT (10 µM), and ciclosporin A (1 µM). NFAT inhibition did not protect the cells from the effects of PMA, and any significant differences were of negligible magnitude (Fig. 3D). Next, we tested different inhibitors of the ERK/MAPK pathways, applying the specific MKK1 (MEK1) inhibitors PD0325901 (100 nM) and PD184352 (1 µM)^74–76^. We also tested the supposed MKK inhibitor PD98059 (1 µM), although its effectiveness is unclear^75,76^. To inhibit p38 MAPK, we used 1 µM BIRB796^74^ (Fig. 3E). The results of this experimental series show that MKK1 inhibition by PD0325901 and PD184352 prevents PMA-induced t-tubule loss, whereas PD98059 and BIRB have very little to no effect. Western blotting showed that PKD phosphorylation was not affected by MKK inhibitors (Fig. S2). This indicates involvement of the ERK1/2 pathway via MKK1 and suggests that p38 MAPK does not contribute. In summary, these results provide strong evidence for the involvement of PKD and MKK1 as a mediator of PMA-induced t-system degradation.

### 3.4. T-Tubule loss is driven by NFκB

Since both PKD and ERK1/2 induce NFκB signaling and show cross-talk in NFκB activation^77^, we next focused on the NFκB pathway, which can be divided into a canonical and a non-canonical pathway. We used a series of NFκB inhibitors that block the pathways at different sites (see Fig. 4A).

**Fig. 4:**
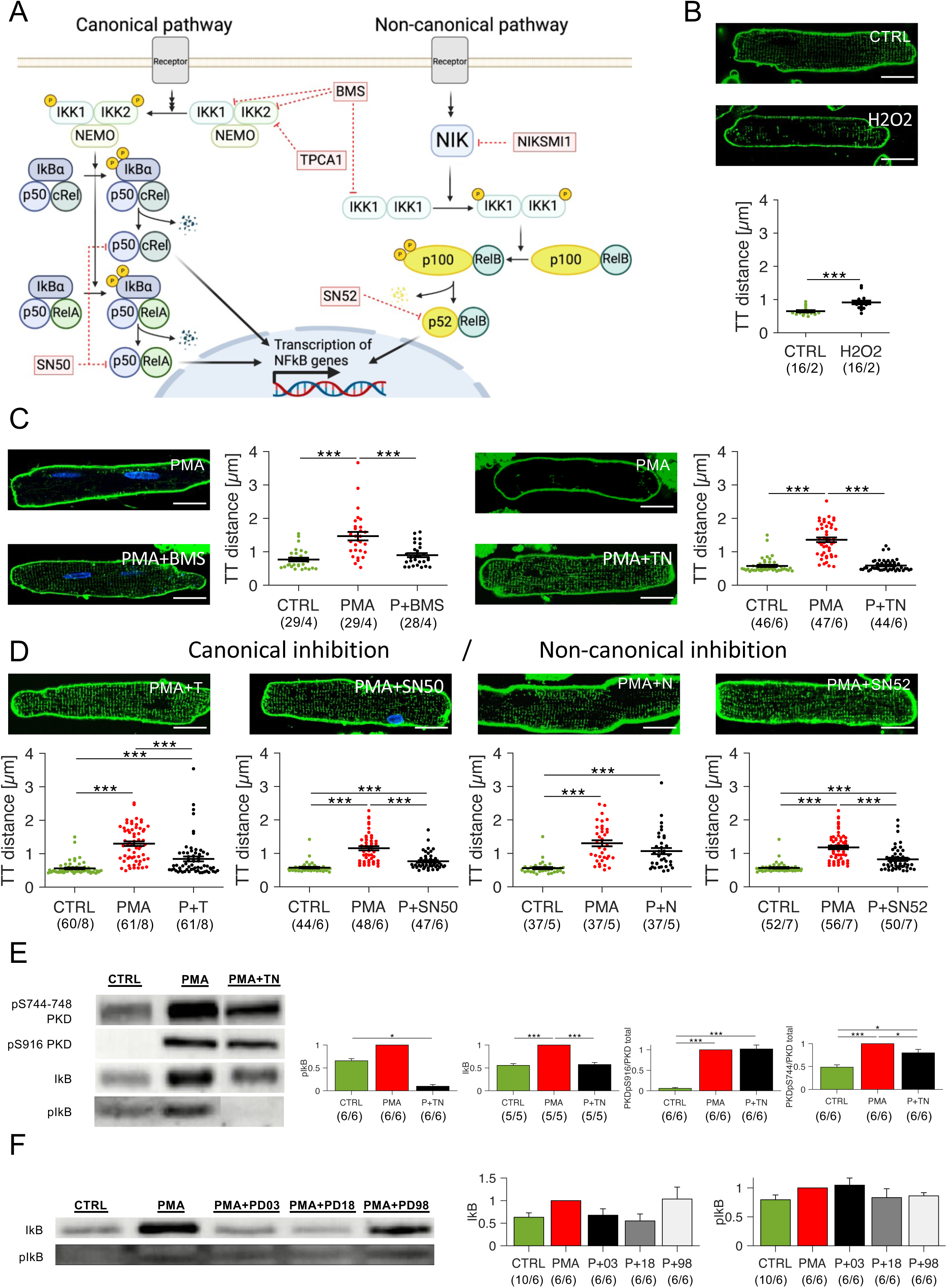
Effects of NFkB inhibition on PMA-induced t-system loss. **(A)** Schematic representation of the pathways which regulate NFkB activity. In the canonical pathway, the NFkB dimers p50-cRel and p50-RelA are bound in an inactive state to IkBa in the cytoplasm; a diversity of signals triggers the phosphorylation and activation of the IKK1-IKK2-NEMO complex, which in turn phosphorylates IkBa, causing its proteasomal degradation. The dimers now translocate to the nucleus and regulate gene transcription. In the non-canonical pathway, the p100-RelB dimer is inactive in the cytoplasm; activation of NFkB-inducing kinase (NIK) phosphorylates and activates the IKK1-IKK1 complex, which triggers the processing of the p100 subunit into its active p52 form. The p52-RelB dimer now translocates to the nucleus and regulates gene transcription. The inhibitors applied in this study are shown in red. BMS blocks IKK1 and IKK2 activity, TPCA1 blocks IKK2 activity, NIKSMI1 blocks NIK activity, SN50 blocks p50-cRel and p50-RelA translocation, SN52 blocks p52-RelB translocation. **(B)** Effect of H_2_O_2_ 50µM on the TATS of rat myocytes after 1 day in culture. Representative confocal images of each group are shown (Di-8-ANEPPS staining). The graph compares TT distance in CTRL and H_2_O_2_-treated cells. **(C)** Effect of inhibiting both NFkB pathways on PMA-induced TATS loss; cells were cultured for 1 h with the inhibitors BMS-345541 20µM (BMS) or TPCA1 10µM + NIKSMI1 500nM (T+N) before PMA addition and then cultured for another 24 h. Vehicle (CTRL) and PMA were served as controls. **(D)** Effect of selective NFkB pathway inhibitors TPCA1 (T), SN50 9µM, NIKSMI1 (N) and SN52 7µM on PMA-induced TATS loss; cells were cultured for 1 h with the inhibitors before PMA addition and then cultured for another 24 h. **(E)** Effect of T+N on PMA-induced PKD and NFkB activation; cells were cultured for 1 h with the inhibitor before PMA addition and then cultured for another 24 h. Example blots can be seen, along with a graph comparing PKD phosphorylation at S744-748 (PKC site) and S916 (PKD autophosphorylation site), normalized to total PKD levels, and IkB and pIkB levels. **(F)** Effect of different MKK1/ERK inhibitors on PMA-induced NFkB activation; cells were cultured for 1 h with the inhibitor before PMA addition and then cultured for another 24 h. NFkB activation was assessed by Western blotting of IkBa. Scale bars: 20 µm; statistics: * p<0.05, ** p<0.01, *** p<0.001, unpaired non-equal variance tests, matched cell isolations. Numbers below graphs indicate the number of cells or data points / the number of cell isolations (hearts).

PMA caused upregulation of IκBa, an NFκB response protein, after 30 min and also after 24 h of treatment (Fig. S3A+B). To activate NFκB without involving PMA or PKC, we added H_2_O_2_^78,79^ at a concentration of 50 µM to rat cardiac myocytes. This led to significant t-tubule loss after one day in culture (*Δ*TT 0.91±0.06 µm vs 0.64±0.03 in CTRL, p<0.001, Fig. 4B), suggesting that NFκB activation independently of PKC induces t-tubule loss as well. Furthermore, we found that NFκB inhibition prevents the spontaneous degradation of the t-system in rat cardiomyocytes cultured for up to three days (Fig. S4). This indicates that the more slowly occurring t-tubule loss which is well known in cultured isolated cardiomyocytes is also driven by NFκB.

We then broadly blocked the NFκB pathway in the presence of PMA, using 20 µM BMS-345541, a substance that inhibits both IKK1 and IKK2^80^ or a combination of 10 µM TPCA1 and 0.5 µM NIKSMI1, which interfere with IKK2 and NIK, respectively^81,82^. Thereby, the canonical and the non-canonical pathways were inhibited (see Fig. 4A). In both cases we found that the PMA-induced t-tubule loss was entirely prevented (Fig. 4C). Western blots showed that PKC and PKD were not affected by these drugs, as visible from unchanged PKD phosphorylation at the PKC site s744-748 and the autophosphorylation site s916 (Fig. 4E, Fig. S3C). IκBa, however, was downregulated significantly in comparison to PMA. This provides evidence for direct blockage of NFκB as a way to preserve t-tubules in the presence of PMA.

To elucidate the contribution of the two major NFκB pathways, we then used TPCA1 or SN50 to exclusively inhibit the canonical pathway, and NIKSMI1 or SN52 to exclusively block the non-canonical pathway. Inhibiting only one of the two pathways resulted in intermediate effects (Fig. 4D), suggesting that both pathways are activated by PMA and contribute additively to t-tubule loss.

Because suppression of NFκB signaling could also underly the effects of the MKK1 inhibitors shown in Fig. 3, we assessed NFκB activation after inhibition of MKK1 by quantifying IκBa via Western blotting (Fig. 4F). Interestingly, our analysis yielded a clearly inhibitory effect of PD03 and PD018 on the expression of IκBa, but not of PD98. This fits to the observation that PD03 and PD018 prevented PMA-induced t-tubule loss, whereas PD98 did not. Combined with the observations that MKK1 inhibitors did not affect PKD (Fig. S2) and NFAT inhibitors had neither influence on t-tubules nor on IκBa (Fig. S3D), these findings further support the hypothesis that NFκB activation underlies t-tubule loss. We could not provide evidence that receptor-mediated PKC activators angiotensin II, endothelin 1 and phenylephrine activate NFκB (Fig. S3E), although this was reported in neonatal cardiomyocytes and smooth muscle cells^83,84^. However, this fits to the relatively small effect size of A+E+P on the t-system when compared with PMA (Figs. 1F and 1B) and may also result from a lower degree of ERK1/2 activation in A+E+P compared to PMA. Collectively, the findings suggest that NFκB activation via both pathways (IKK1, IKK2) as well as via the ERK1/2 pathway is accountable for maximal acceleration of t-tubule loss.

### 3.5. RNA Sequencing supports NFκB signaling

ERK1/2 can activate NFκB via phosphorylation of p65 by mitogen- and stress-activated protein kinase-1 (MSK1)^85^ and IKK2 may activate ERK^77^. Our results indicate that inhibition of MKK1, an upstream activator of ERK1/2, prevents NFκB activation and t-tubule loss in rat cardiomyocytes (Fig. 4F). Although this is still consistent with NFκB being the main signal causing t-tubule loss, it raises the question whether NFκB is upstream of ERK1/2 and the actual effector is ERK pathway activation. To investigate those possibilities and to explore other targets that could be responsible for t-tubule loss, we performed RNA sequencing in rat myocytes isolated from six hearts. Cells from each heart were divided into three groups and treated for a total of 5 h with either vehicle as control, PMA or NIKSMI1+TPCA1+PMA (PMA+TN). The PCA-based cluster analysis (Fig. 5A) revealed a clear effect of the treatments, especially of PMA, but also of the biological replicate (cell isolation or batch effect). Therefore, the statistical analysis for the comparison of treatments was corrected for the effects of the cell isolation by adding the heart ID as a covariate. Volcano plots (Fig. 5B) showed a large amount of differentially expressed genes (DEGs, p < 0.05) in PMA vs CTRL (up: 2546, down: 2573) and a comparably small amount in PMA+TN vs PMA (up: 632, down: 629). Therefore, regarding the number of DEGs, PMA+TN vs CTRL was closer to PMA vs CTRL, with a total of 4803 DEGs. This indicates that PMA regulated a large number of genes via different pathways, as the effect of NFκB inhibition by NIKSMI1+TPCA1 on gene expression was much more restricted. See the supplemental data for the full list of DEGs.

**Fig. 5:**
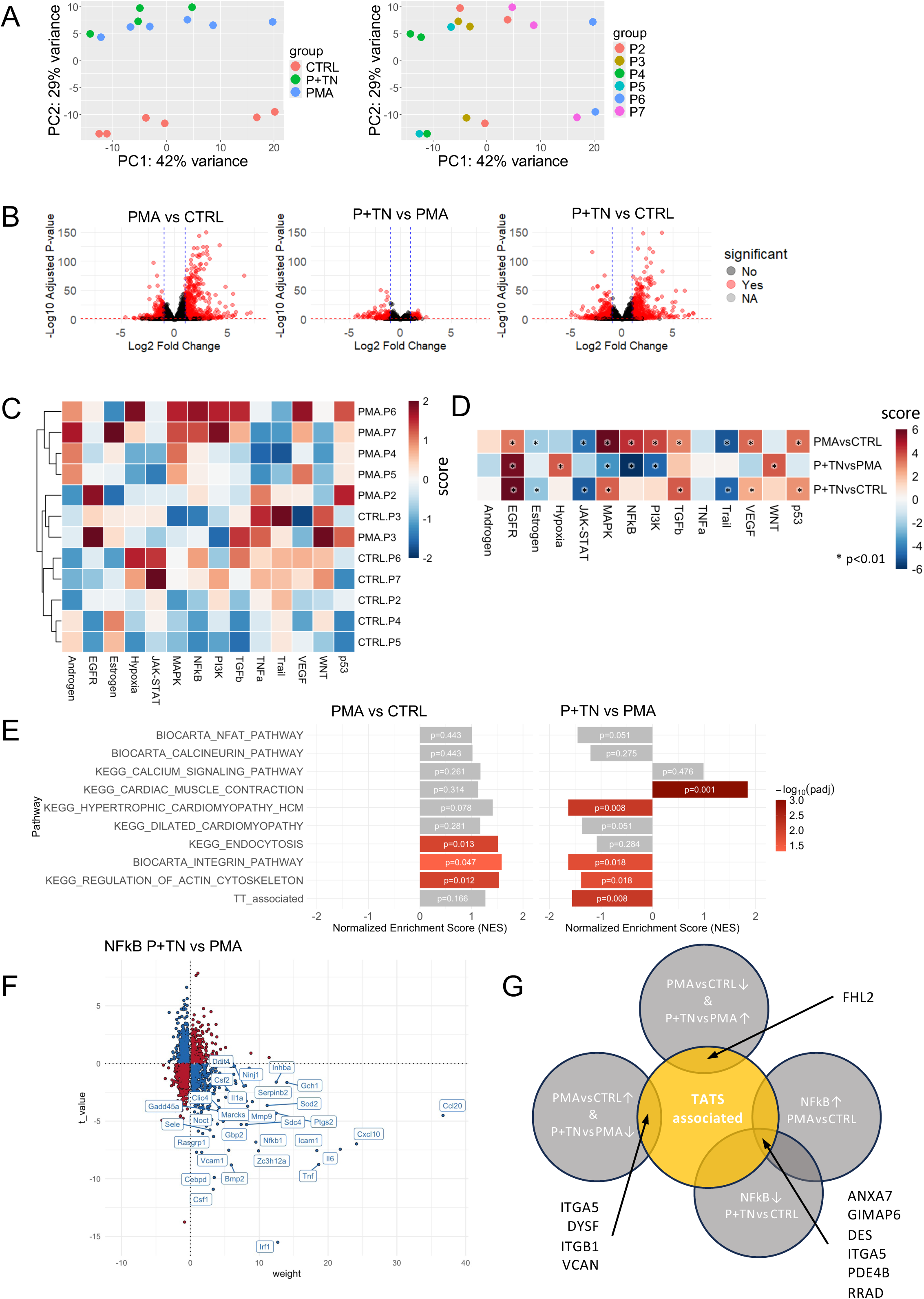
Gene expression analysis by RNASeq in isolated rat cardiomyocytes. **(A)** Principal component analysis (PCA) of individual samples. Colors indicate treatment (control (CTRL), PMA and NIKSMI1+TPCA1+PMA (P+TN)). Or the heart, from which the cells were isolated (P2-P7). **(B)** Volcano plots of differentially expressed genes after calculation of the log_2_-fold change and the adjusted p value (Benjamin-Hochberg correction) for each gene in PMA vs CTRL, P+TN vs PMA and P+TN vs CTRL, taking batch effects (hearts) as covariate into account. The significance threshold was set to the adjusted p<0.05. **(C)** Heatmap showing the score of a PROGENy pathway analysis in individual PMA and CTRL samples, using the log-normalized counts. The lines of the tree on the left indicate the Euclidean distance between the samples. The score was obtained by scaling column-wise (subtraction of mean, followed by division by the standard deviation). **(D)** Same as (C) but using the Wald statistics value of the differential expression analysis of PMA vs CTRL, P+TN vs PMA and P+TN vs CTRL. **(E)** Gene set enrichment analysis (GSEA) of PMA vs CTRL and P+TN vs PMA. The direction and magnitude indicate the normalized enrichment score, while the color indicates the adjusted p value. **(F)** Genes with their weights from the PROGENy NFkB pathway model and their t_value from the multilinear model. Genes contributing to the downregulation of the pathway are shown as blue circles, genes contributing to upregulation as red circles. The 30 genes with the highest score (absolute value of weight multiplied with t_value) are labelled, i.e. the genes contributing most to downregulation of the NFkB pathway in P+TNvsPMA. **(G)** Schematic showing the approach to explore genes regulated by PMA and NFkB that might contribute to TATS loss or maintenance. The orange circle indicates genes that were associated with the TATS in previous studies. The other circles indicate genes identified after differential expression analysis oppositely regulated in PMAvsCTRL and P+TNvsPMA or those contributing to NFkB pathway upregulation in PMAvsCTRL and downregulation in P+TNvsCTRL. The intersections of the resulting lists with the TATS-associated gene list are indicated by arrows and the resulting genes are given next to the arrows.

To investigate the effects of treatments on pathway activation, we then performed a PROGENy pathway analysis^48^, using the log-transformed normalized counts per sample (Fig. 5C). Comparing PMA with control, the results show clustering of the groups, with upregulated EGFR, p53, Androgen, VEGF, MAPK, PI3K and NFκB pathways and predominantly downregulated JAK-STAT, WNT and Trail pathways in PMA. A heatmap showing all samples, including PMA+TN, can be retrieved from Fig. S5A. At close inspection, the effects of the cell isolation (heart), indicated by P2 through P7, become visible also in this representation. Therefore, in Fig. 5D, we performed an analysis using the Wald statistic (log_2_-fold change divided by its SE) from the differential expression analysis of the groups against each other as input parameter (PMA vs CTRL, PMA+TN vs PMA and PMA+TN vs CTRL). This again compensated for batch effects (hearts). We found that PMA significantly upregulated the MAPK and NFκB pathways. As expected, the addition of NIKSMI1+TPCA1 had a strong inhibiting effect on the NFκB pathway, but – importantly – only mildly affected the MAPK pathway, as seen from the significant upregulation of MAPK in PMA+TN vs CTRL. This allows the conclusion that t-system preservation by NFκB inhibitors was not mediated by blocking the MAPK pathway. Some other pathways, e.g. p53 or VEGF, showed similar patterns and are therefore unlikely to be involved in t-tubule loss or maintenance. The EGFR and JAK/STAT pathway alterations were even enhanced by NIKSMI1+TPCA1 and can be excluded as well (see Fig. S5B-D and Table S1 for more details). The PI3K-AKT pathway was activated by PMA and reversed to control levels by NIKSMI1+TPCA, leaving it as a candidate besides NFκB. However, inhibition of AKT by two reliable inhibitors did not rescue the t-system (Fig. S5G), ruling out the PI3K-AKT pathway. Altogether, these findings largely excluded pathways other than NFκB and render it unlikely that signals downstream of ERK1/2 other than NFκB are involved in t-tubule loss, even though ERK1/2 may contribute to NFκB activation or modulate the NFκB effects.

We then performed a gene set enrichment analysis (GSEA) to investigate the NFAT pathway, which is not included in the PROGENy database, and to investigate gene sets related to Ca^2+^ signaling, cardiac diseases and function as well as t-tubule associated genes^51^ and pathways (Fig. 5E). The two chosen gene sets representative of NFAT activation (NFAT and calcineurin) were not significantly enriched in PMA vs CTRL nor in PMA+TN vs PMA, providing no evidence for a contribution of NFAT signaling to t-system breakdown. This is consistent with Fig 3D, where NFAT/calcineurin inhibitors were ineffective. The gene sets associated with Ca^2+^ signaling, cardiomyopathy and t-tubules were also not significantly enriched in PMA. Interestingly, endocytosis-, integrin- and cytoskeleton-related genes were significantly upregulated. In PMA+TN vs PMA, however, genes related to cardiac muscle contraction were upregulated, hypertrophic cardiomyopathy related genes were downregulated and t-tubule associated genes were downregulated, as well as genes related to the integrin pathway and the cytoskeleton. This suggests that downregulation of some t-tubule related genes may stabilize the t-system or that t-tubule loss by PMA induces upregulation of these genes via NFΚB. However, indirect or compensatory effects may also underlie these findings.

To further explore the genes that could be responsible for the loss of t-tubules, we investigated the genes contributing to NFκB downregulation in PMA+TN vs PMA, i.e. those genes that have a positive weight and were downregulated or have a negative weight and were upregulated (blue dots in Fig. 5F). It seems likely that among these genes one could find genes responsible for t-tubule loss or preservation. The 30 genes with highest weight and change are labelled. However, because the number of candidate genes was still enormous, we additionally restricted them as follows: We selected those genes in the PROGENy NFκB pathway that were 1) upregulated by PMA vs CTRL and had a positive weight or downregulated and had a negative weight, 2) downregulated in PMA+TN vs PMA and had a positive weight or upregulated and had a negative weight, and 3) were contained in the list of t-tubule associated genes (Fig. 5G). This resulted in six genes: ANXA7, a membrane binding protein, GIMAP6, a protein involved in autophagy and immune cell function, DES, an intermediate filament, ITGA5, an integrin, PDE4B, a phosphodiesterase, and RRAD, a GTP-binding protein. When creating the intersection of genes significantly upregulated by PMA vs CTRL and downregulated by PMA+TN vs PMA or vice versa and t-tubule associated genes, we found again ITGA5, but also DYSF, ITGB1, VCAN and FHL2. In sum, these genes appear to be related to membrane traffic, repair and anchoring in the extracellular matrix as well as mutations with cardiac phenotype^86,87^, which would fit well to a possible role in t-system degradation and maintenance.

### 3.6. NFκB activation by PMA affects excitation-contraction coupling

The t-system opposes L-type Ca^2+^ channels (LTCC) in the t-tubular membrane to the ryanodine receptors (RyRs) in the sarcoplasmic reticulum (SR) and is therefore important for cardiomyocyte cytosolic Ca^2+^ release. We investigated how t-tubule loss by PMA and its prevention by NFκB inhibition affect the density and mutual distance of RyRs and LTCCs and thereby myocyte function (Fig. 6). Using 3D confocal microscopy and automated image analysis, we found that PMA did not change the RyR cluster density of rat myocytes after 24 h of treatment (Fig. 6A). LTCC density, however, was lower in PMA than control cells and increased in PMA+TN versus PMA, although PMA+TN did not reach control level (Fig. 6B). As a result, the distance between RyR clusters and their closest LTCC cluster (Fig. 6C) was significantly higher in PMA-treated cells (1.02±0.03 µm) than in CTRL (0.76±0.02 µm) and PMA+TN (0.91±0.02 µm). The cells were also co-stained for JPH2, which showed similar changes as LTCC (Fig. S6A). BIN1 density was slightly reduced and more irregular in PMA, but completely preserved in P+TN (Fig. S6B). We tested the functional effects of reduced t-system density and increased RyR-LTCC distances by dual imaging of the Ca^2+^ indicator Fura-2 and the sarcomere length. Fig. 6D displays representative examples of the Ca^2+^ transients at 0.5 Hz pacing rate. It is evident that PMA treatment caused smaller amplitudes and this effect was prevented by NFκB inhibition via TN (Fig. 6E). Similarly, sarcomere shortening, an indicator of contractility, was significantly reduced in PMA, but preserved in P+TN (Fig. 6F). Comparable effects were achieved by PKD inhibition (Fig. S6C). The loss of t-tubules decreases the efficiency by which Ca^2+^ release from the SR is triggered. One would therefore expect a decrease in the fraction of Ca^2+^ release from the SR. By application of caffeine, which causes full release of the SR Ca^2+^ it is possible to estimate the fractional SR Ca^2+^ release (Fig. 6G). It becomes clear from the examples that not only the fraction, but also the amount of Ca^2+^ inside the SR was decreased in PMA. This was confirmed by quantification and statistical analysis (Fig. 6H). Interestingly, the decay time was longer in PMA, suggesting a reduced Ca^2+^ extrusion via the Na^+^-Ca^2+^-exchanger^88^. To confirm that the observed effects were due mainly to t-tubule loss, we additionally performed confocal line scans (Fig. S6D). Line scans show that Ca^2+^ release was delayed and diminished in regions inside the cell with high TT distance, which were more abundant in PMA treated cells than in CTRL and P+TN cells. In summary, these results provide strong evidence for impaired excitation-contraction coupling in PMA-treated cells due to t-system degradation, which could be prevented by NFκB inhibition.

**Fig. 6.**
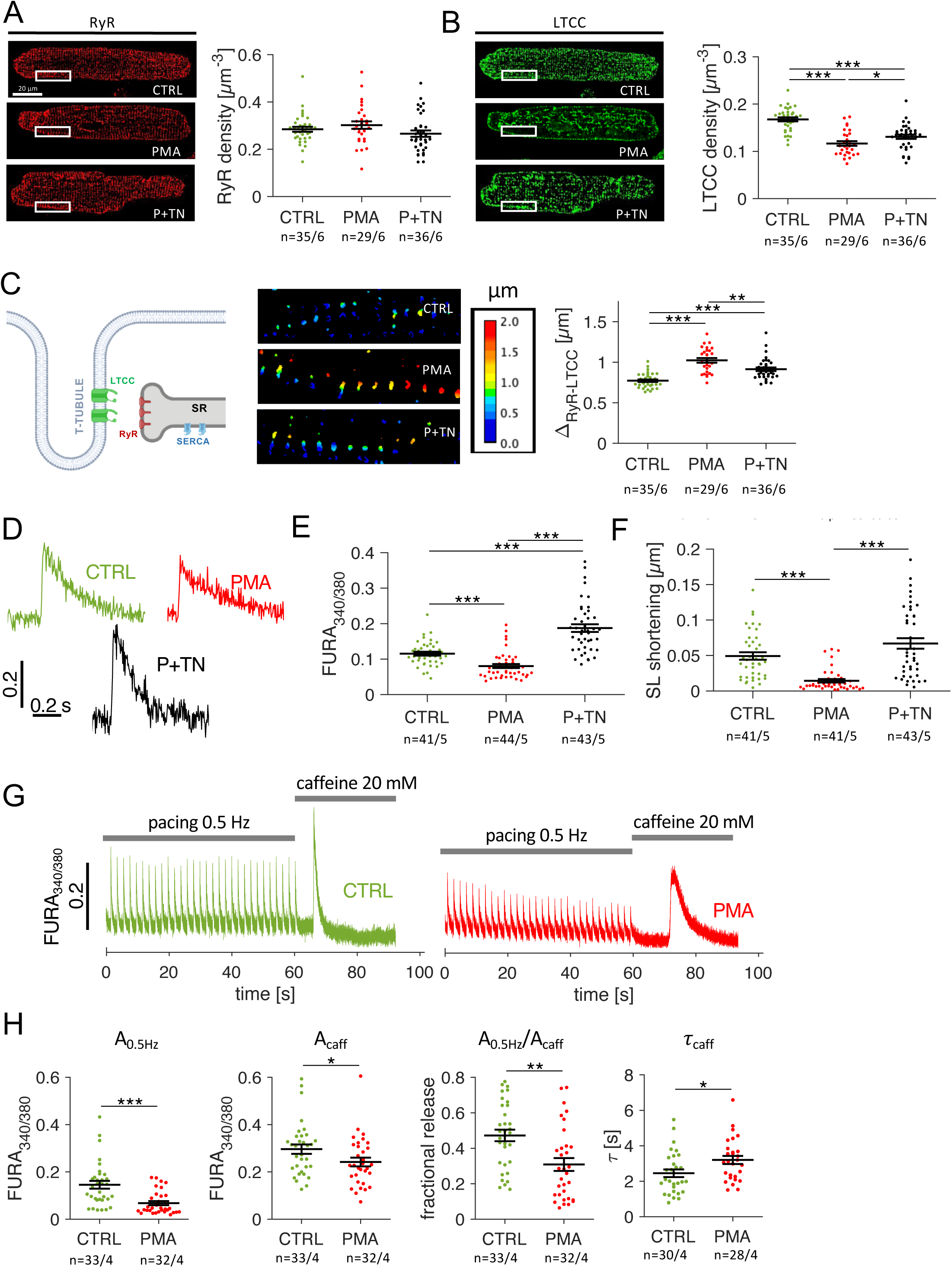
Effects of PMA and NFkB inhibition on excitation-contraction coupling. **(A)** Isolated rat myocytes treated with vehicle (CTRL), PMA or PMA + NFkB inhibitors TPCA1 + NIKSMI1 (P+TN) for 24 where stained by immunofluorescence for ryanodine receptors (RyRs) and imaged by 3D confocal microscopy. RyR density was calculated as the fraction of voxels occupied by RyR signal within the cell. **(B)** Same cells as shown in A, co-stained for L-type Ca^2+^ channels (LTCC) with quantification of LTCC density. **(C)** The distance between RyR clusters and their closest LTCC cluster was calculated by creating the LTCC distance map. Shown are magnified views of the regions indicated by white boxes in (A) and (B). The average distance of RyR-positive voxels was then used to obtain the RyR-LTCC distance (ι1). **(D)** Example traces of the Ca^2+^ signal of Fura-2-loaded cells at 1 Hz pacing rate at 35° C. **(E)** Amplitude of the Ca^2+^ transients. **(F)** Sarcomere length (SL) shortening, i.e. absolute difference to resting length, during contraction. **(G)** Example traces of a CTRL and a PMA treated cell with 0.5 Hz pacing for 60 s and a subsequent switch to caffeine superfusion to elicit complete SR Ca^2+^ release. Pacing was stopped during caffeine superfusion. **(H)** Ca^2+^ signal amplitude at 0.5 Hz pacing (A_0.5Hz_), during caffeine superfusion (A_caff_), fractional release (=A_0.5Hz_/A_caff_) and the time constant of the Ca^2+^ signal decay after the caffeine-induced release (τ_caff_).

### 3.7. Mechanism of t-tubule loss

To study the fate of lost t-tubules, we conducted experiments with fluorescent dextran to track endocytosis-related internalization of membrane components. Anionic dextrans are hydrophilic polysaccharides that cannot permeate intact membranes. Thus, unless the plasma membrane integrity is compromised^45,8^, they can enter the cytosol only by active processes like endocytosis. This results in intracellular compartments or vesicles filled with dextran^89,90^. Fig. 7A illustrates our methodological approach to test the hypothesis that t-tubules disappear by internalization. The example image of a rat cardiomyocyte (Fig. 7B) shows dextran components inside the cell that are structurally very similar to t-tubules. Notably, structures positive for dextran were not stained by AM4-66, indicating that dextran is entrapped in membranes not connected to the surface membrane (magnification in Fig. 7B and Fig. S7A). We quantified the overlap of both signals and found in CTRL cells that only 11.8% ± 0.5% (n = 287/40) of all dextran-positive voxels were also positive for AM4-66. In a positive control experiment, we incubated dextran and AM4-66 simultaneously (Fig S7B). Here, we found a much higher overlap of 47.6% ± 1.4% (n = 30/2). Note that a 100% overlap was not to be expected, as AM4-66 stains the membrane and dextran the contents. Importantly, this confirms that the dextran-components are surrounded by membranes that were previously connected to the cell membrane and points towards an endocytic process. The mutual exclusion of the signals when stained sequentially, suggests that the internalized t-tubule-shaped structures are disconnected from the plasma membrane. If the t-tubules had remained connected to the plasma membrane, the lipophilic dye AM4-66 would have also stained the dextran-surrounding membrane.

**Fig. 7.**
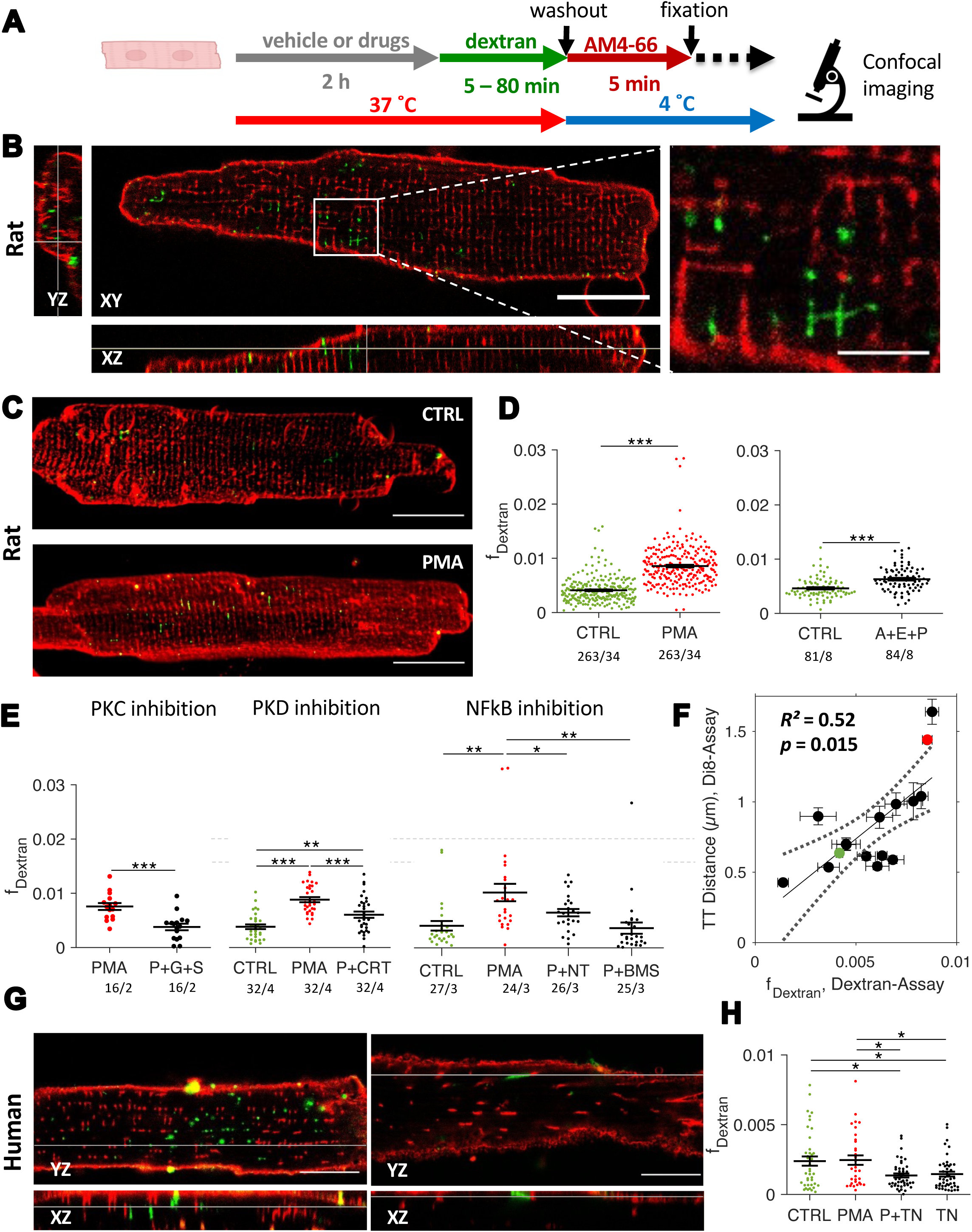
Internalization of t-tubules and modulation of the ratio with activators and inhibitors. **(A)** Methodological approach. **(B)** 3D confocal image showing the xy, xz and xz planes of an isolated rat cardiomyocyte incubated with dextran for 20 minutes (green) and stained with the membrane dye AM4-66 (red). A magnified view of the white box is shown on the right. **(C)** 3D projection of a 10 µm stack of an isolated cardiomyocyte incubated with PMA or vehicle (CTRL) for 2 h. Internalized dextran (green) and AM4-66 staining (red) are shown. **(D)** Quantification of the volume ratio (fraction) of internalized dextran (f_dextran_) of control rat cells (CTRL) and rat cells treated for 2 h with PMA or angiotensin-II + endothelin-1 + phenylephrine (A+E+P). **(E)** f**_dextran_** of cells treated with PMA and the PKC inhibitors GÖ6893 (G) and sotrastaurin (S), PKD inhibitor CRT0066101 (CRT), and NFκB inhibitors TPCA1 (T) + NIKSMI1 (N), and BMS345541 (BMS). **(F)** Correlation of f_dextran_ in cells incubated with different pharmacological compounds for 2 h with the respective average of the mean t-tubular (TT) distance of cells stained with Di-8-ANEPPS after treatment for 1 day. The PMA group is depicted in red, the CTRL group in green. **(G)** 3D confocal images showing the xy and xz planes of isolated human heart failure cardiomyocytes with internalized dextran (green) and AM4-66 membrane staining (red). The left image shows a cell with a dense TATS, the right image a cell with pathological remodeling. **(H)** f**_dextran_** of human cells treated with PMA and the NFκB inhibitors TPCA1 (T) + NIKSMI1 (N). Analysis Statistical test: paired t-test. P-values after correction for multiple comparisons: *p < 0.05; **p < 0.01; ***p < 0.001. Scale bars: 20 µm, magnification in (A) 5 µm.

Given these results, we next asked whether PKC/NFκB-induced t-tubule loss occurs via this mechanism. After 2 h of PMA treatment, we observed significantly higher amounts of intracellular dextran than in control, as visible from the 3D projection shown in Fig. 7C and the 2-fold increased fraction of internalized dextran in PMA (Fig. 7D). This resulted from an increased number and size of dextran-positive intracellular components (Fig. S7C). As receptor-mediated PKC activation caused t-tubule loss as well (Fig. 1F), we tested the combination of A+E+P and found that they significantly increased the amount of intracellular dextran (Fig. 7D). The extent was not as great as with PMA, which fits well with the comparatively lower degree of t-tubule loss, suggesting a direct link between the rate of dextran internalization and t-tubule loss. To further test the link between dextran uptake and t-tubule loss in rat cells, we used inhibitors of PKC (GÖ6983 and sotrastaurin), PKD (CRT0066101) and NFκB (BMS and TPCA1+NIKSMI), which all were able to significantly prevent PMA-induced t-system degradation (Figs. 2-4). Here we found that they also significantly decreased the fraction of internalized dextran (Fig. 7E). In total we tested 16 compounds (Supplemental Table 2). In Fig. 7F, we plotted the average fraction of dextran in all imaged cardiomyocytes against the mean t-tubule distance after 24 h of treatment. Linear regression revealed a positive and significant correlation (p = 0.015, R² = 0.52). Collectively, these results provide evidence that the internalized dextran corresponds to internalized t-tubules, allowing the conclusion that t-tubule loss in isolated cardiac myocytes occurs via an endocytic process.

We next examined if the same observations could be made in human cardiomyocytes obtained from failing hearts, which often display a remodeled t-system. Fig. 7G presents 3D confocal images of a cell with a dense t-system and another one with t-system remodeling, including reduced density and sheet-like changes in t-tubule shape^7^. It is clearly visible that in both cases the internalized dextran fits to the shape of the respective t-tubules, i.e., tubular in the cell with a dense t-system and sheet-like in the cell with remodeled t-system. Application of PMA for 2 h did not increase the amount of dextran uptake in human cells. However, NFκB inhibition by TPCA1+NIKSMI1 significantly reduced the internalized dextran both vs CTRL and vs PMA (Fig. 7H), suggesting that also in human cells NFκB activation is involved in this process.

### 3.8. T-tubule internalization via a macropinocytosis-related process

After identifying t-tubule internalization as a mechanism, the next step was to study endocytosis-related processes as potential underlying factors. We tested various endocytosis inhibitors in combination with PMA and evaluated the fraction of internalized dextran compared with PMA alone. Dynasore, an inhibitor of dynamin and clathrin-mediated endocytosis^91^, had only little effect. In contrast, pitstop 2, which potently inhibits clathrin-dependent and clathrin-independent endocytosis^92^ and disrupts actin dynamics^93^, was very effective (Fig. S8A). This is compatible with macropinocytosis, which relies required actin polymerization. Macropinocytosis also fits well to the large size of internalized membrane. Several macropinocytosis inhibitors showed a strong and significant effect in reducing the internalization ratio (Fig. 8A). Amiloride, which inhibits the Na^+^/H^+^ exchanger, impairs actin polymerization at high concentrations (2 mM) by causing acidification of the submembranous cytosol^94^. Amiloride reduced the intracellular dextran fraction significantly when compared with PMA, even though it did not fully reach control levels. Next, we tested the PI3K inhibitor wortmannin. PI3K is necessary for the initiation of actin polymerization^95–97^ and involved in many processes of endocytosis and membrane traffic^98^. Similar as amiloride, wortmannin decreased the amount of internalized dextran. This was replicated with PI103, another PI3K class I inhibitor (Fig. S8B). We also confirmed that PI3K inhibition did not affect NFκB signaling (Fig. S8C), but it prevented the PMA-induced loss of the t-system after 1 d (Fig. S8D). Since actin polymerization is disturbed by PI3K inhibition and is required for endocytosis and macropinocytosis, we used cytochalasin D^99^ as a direct inhibitor of actin polymerization. This compound also reduced the fraction of dextran in comparison to PMA. Cytochalasin D also prevented the PMA-induced degradation of the t-system after 1 d (Fig. S8E). We also tested two FDA-approved macropinocytosis inhibitors^100^, auranofin and imipramine. Both had a significant, but intermediate effect in reducing the PMA-induced dextran internalization (Fig. S8D+E). Because myosin 1 is required for macropinocytosis^101,102^, we next tested the effects of a myosin 1 inhibitor, pentachloropseudilin (Fig. 8B). Pentachloropseudilin completely inhibited the PMA-induced uptake of dextran into the cells. Inhibition of myosin 2 by blebbistatin, myosin 5 by pentabromopseudilin and myosin 6 by triiodophenol did not prevent PMA-induced t-system loss(Fig. S8G).

**Fig. 8:**
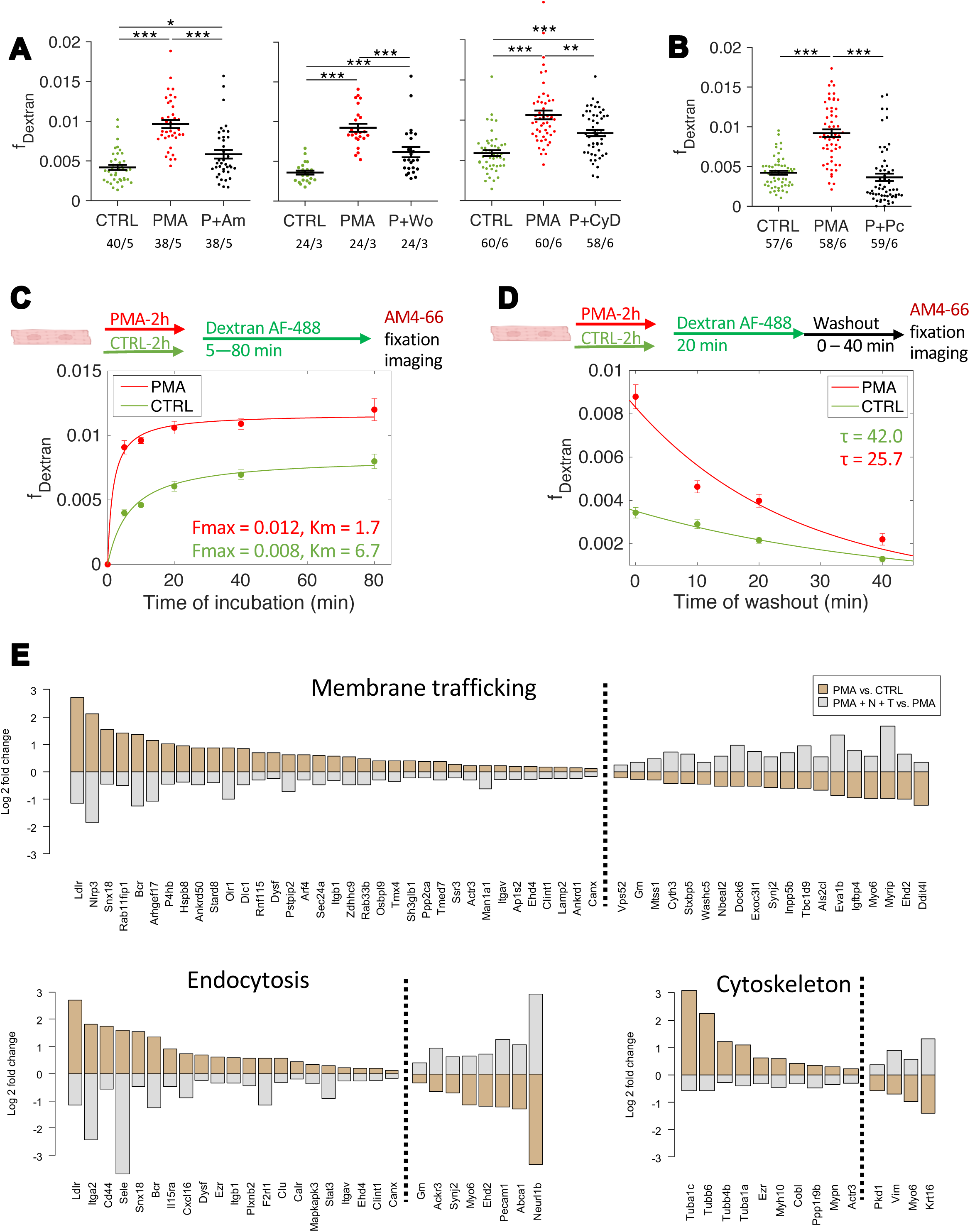
Mechanisms of t-tubule internalization. **(A)** Quantification of the volume ratio of internalized dextran in cardiomyocytes incubated with PMA and various macropinocytosis inhibitors: amiloride (Am), wortmannin (Wo), and cytochalasin D (CyD). **(B)** Quantification of internalized dextran in the presence of the myosin I inhibitor pentachloropseudilin (Pc). Statistical test: paired t-test. P-values after correction for multiple comparisons: *p < 0.05; **p < 0.01; ***p < 0.001. **(C + D)** Methodological approach for experiments with different incubation and washout times of dextran 10 kDa AF-488. Cardiomyocytes were incubated with PMA (50 nM, red) or vehicle (CTRL, green) for 2 hours, followed by dextran application for 5–80 minutes **(C)** or dextran application for 10 minutes followed by a washout for 10–40 minutes **(D)**. The fraction of intracellular dextran in myocytes at different time points was quantified and visualized as the mean ± standard error. In **(C)**, data were fitted using an uptake curve, following saturation kinetics, and in **(D)**, a negative exponential fit was applied. **(E)** Results of RNA sequencing (RNA-seq): Differentially expressed genes (adjusted p-value < 0.05) that were upregulated by PMA and downregulated by the combination of PMA with NIKSMI + TPCA1, and vice versa. The log2 fold-change of genes associated with vesicle trafficking, endocytosis, and cytoskeletal proteins is shown.

Next, we assessed the uptake and degradation rates of internalized dextran by varying the incubation and washout times. Specifically, we incubated rat cardiomyocytes for 2 h with either PMA or vehicle (control), followed by dextran incubation for 5 to 80 min and then fixed the cells, or we incubated them for 20 min and fixed them later after varying washout times up to 40 min in culture medium. When varying the incubation time (Fig. 8C), we observed a quick increase in the fraction of internalized dextran, which then slowed down. The data could be fitted well by saturation kinetics, suggesting a process with limited rate and saturation, representable by an uptake curve. The maximum fraction of dextran was 0.008 in CTRL and 0.012 in the PMA-treated cells. The time at half-maximum (K_m_) was 6.7 min for CTRL and 1.7 min for PMA, suggesting that PMA treatment increased both the rate and capacity of dextran uptake, perhaps by increasing the number or efficiency of macropinocytic events. The saturation also suggests degradation or removal of dextran. When evaluating different washout times (Fig. 8D), both CTRL and PMA-treated cells showed a significant loss of dextran over time. In an exponential decay model, the time constant (*τ*) in CTRL was 42.0 min, while in PMA *τ* equalled 25.7 min. This indicates that PMA-treated cells cleared the internalized dextran more quickly than CTRL and that the higher saturation value in PMA was due to faster uptake rates. This suggests that PMA enhances not only the uptake but also the degradation or recycling of internalized components, potentially altering the dynamics of t-tubule membrane turnover.

As these results point towards important roles of membrane traffic, endocytosis and cytoskeletal dynamics in t-tubule loss, Fig. 8E presents differentially expressed genes from the RNA sequencing related to these ontologies in more detail. We focused on differentially expressed genes that were oppositely regulated when treated with the PKC activator PMA compared to control, and with NFκB inhibitors in combination with PMA compared to PMA alone. In addition to the results presented in Fig. 5F+G, this collection may also contain genes important for t-tubule loss and preservation, as they are related to the identified mechanism and influenced by the treatments that accelerate and prevent t-system disintegration. Taken together these findings indicate that t-tubule loss results from macropinocytic events that lead to membrane internalization and degradation and can be modulated by PKC via NFκB.

## 4. Discussion

Loss and remodeling of the cardiomyocyte t-system are widely recognized as major contributors to electromechanical dysfunction in heart disease – particularly in heart failure with reduced ejection fraction (HFrEF)^103^ – making the t-system a potential target for therapeutic intervention. However, not least due to our lack of understanding of the cellular signals and mechanisms that promote t-tubule loss, the development of targeted therapies remains an unmet challenge. As the disturbances of cellular signals and functions in heart failure are highly complex and closely intertwined with t-system remodeling, investigating the relevant pathways and mechanisms in vivo is particularly difficult^104^. Here, we demonstrate that sustained PKC activation by the phorbol ester PMA causes substantial t-tubule loss in rat, rabbit and human cardiomyocytes through the PKC-PKD-NFκB signaling axis. Mechanistically, we discovered that t-system membranes undergo internalization in a manner compatible with macropinocytosis and that PKC-NFκB activation increases the rate of internalization, while NFκB inhibition reduces it. We also show that inhibiting NFκB rescues excitation-contraction coupling by preserving the t-system. These findings expand our understanding of how t-tubules are actively degraded in cardiomyocytes, placing NFκB at the center of pathological membrane remodeling.

### 4.1. Overall Significance of PKC-Mediated T-Tubule Remodeling

Although little is known about PKC-driven t-tubule loss or remodeling, one study reported that transient PKC activation – particularly via PKCα/β - induces actin-cytoskeletal changes that subsequently damage t-tubules in mouse cardiac myocytes^17^. Our findings both complement and extend these findings. While the earlier study focused on immediate cytoskeletal rearrangements following transient PKC activation, our data reveal that sustained PKC activation leads to t-tubule loss through downstream PKD-NFκB signaling that involves gene expression (Fig. S9). Importantly, we validated key aspects of this pathway in human myocytes and a tissue culture model. In agreement with previous work^17,105^, we confirm that disrupting cytoskeletal actin assembly, e.g. with cytochalasin D, prevents t-system degradation. However, our results suggest a new interpretation: an actin-dependent form of endocytosis, resembling macropinocytosis, is responsible for the internalization and subsequent degradation of t-tubule membranes. This process is upregulated via the PKC-NFκB axis and can be blocked by inhibitors of actin assembly and macropinocytosis (Fig. 8A).

Receptor-mediated activation of PKC by the combination of angiotensin II, endothelin-1, and phenylephrine (A+E+P) also induced significant t-tubule loss, although its effect was less pronounced than that observed with PMA. Notably, our data (Fig. S1C+D) reveal that A+E+P treatment leads to a strong PKD S916 phosphorylation at 2 h, which markedly declines by 24 h. In contrast, PMA maintains robust PKD S916 phosphorylation for 24 h (Fig. S3), indicating that PMA produces a more sustained activation of the PKD-NFκB pathway. This may reflect the differences between receptor-mediated signaling and the direct, potent activation provided by PMA. The reduction in receptor-induced PKD activation after 24 h may be attributable to negative feedback mechanisms or receptor desensitization. It is also possible that A+E+P is degraded more quickly than PMA. In any case, these results suggests that the duration and intensity of PKC-PKD-NFκB signaling are critical determinants of the remodeling process in cardiomyocytes.

### 4.2. Role of Different PKC Isoforms and PKD

PMA activates both classical and novel PKC isoforms^106^, of which the novel isoforms (*δ*,*ε*,*η*,*θ*) are supposed to activate PKD^107^. Guo et al^17^ suggested that the classical isoforms *α* and *β* mediate PMA-induced t-system deterioration in mouse cardiomyocytes, based on in-vivo experiments using enzastaurin administration and in-vitro overexpression of mutant PKC*α* variants. However, despite testing both enzastaurin and ruboxistaurin as PKC*β* inhibitors (Fig. 2), we were unable to fully reproduce these findings. This discrepancy may arise from off-target effects of enzastaurin, e.g. the inhibition of additional kinases and isoforms^108^, or indirect in-vivo effects. It is also conceivable that PKD or NFκB signaling in mouse cardiomyocytes relies more on alternative activation pathways. For instance, some earlier studies have noted facilitation of IKK phosphorylation by PKC*α*^109^. In our study, however, the data strongly indicate that PKD activation by PKC is essential for t-tubule loss because the specific blocker CRT0066101^72^, which selectively blocked the autophosphorylation site of PKD without affecting its PKC-induced phosphorylation site, was highly effective in preventing PMA-induced t-system degradation – which was the case as well with KB-NB142-70^73^ (Fig. 3B+C). This leads us to propose that novel PKC isoforms, such as PKC*δ* or PKC*ε*, are primarily responsible for triggering t-tubule loss by PMA. Some studies implicate PKC*ε* as the central mediator, especially following G_q_-coupled receptor activation in cardiomyocytes^110,111^, while others favour PKCδ-mediated PKD activation^112^. However, the lack of highly specific inhibitors for individual PKC isoforms^76^, necessitates further investigation using alternative approaches (e. g., overexpression of mutant isoforms) in future studies.

### 4.3. NFκB as a Crucial Downstream Effector

PKC and PKD have many downstream pathways, of which we identified NFκB as the most important driver of t-tubule degradation. In accordance with this, all compounds that inhibited NFκB also prevented the loss of t-tubules. As glucocorticoids also inhibit NFκB, this fits to a previous study in which we showed that dexamethasone stabilizes the t-system in cell culture^15^. Blocking only the canonical or the non-canonical NFκB pathway resulted in intermediate effects, indicating that both pathways contribute additively. This also supports a correlation between the intensity and duration of NFκB activation and the degree of t-system damage. Intriguingly, our findings point towards cross-activation through the MKK1-ERK1/2 pathway. ERK1/2 can activate NFκB via phosphorylation of p65 by mitogen- and stress-activated protein kinase-1 (MSK1), as reported in non-muscle cells^85^. The ERK1/2 pathway has also been reported to activate NFκB, by translocation^113^ or by transactivation^114^. Based on our findings, we suggest that ERK1/2 may constitute an important amplifier or modulator of the PKD-NFκB axis in cardiomyocytes and that multiple inputs may converge on NFκB to drive membrane remodeling. This could be an additional explanation for the less pronounced effects of G_q_-protein coupled receptor agonists on the t-system when compared with PMA, as PMA may be a stronger activator of the MKK-ERK1/2 pathway. The requirement for joint activation of PKD and ERK1/2 and the detailed steps leading to full NFκB activation in adult cardiac myocytes may be of great interest not only from a cell biology, but also from a clinical perspective and therefore seem worth further research. We also found that NFκB activation is responsible for spontaneous degradation of the t-system in isolated cardiac myocytes, as NFκB inhibition improved t-tubule density over control (Fig. S4). This may be explained activation of PKD and NFκB after the loss of cell-cell and cell-matrix contacts^115^.

### 4.4. Transcriptomic Perspective and Potential Molecular Targets

Consisting of a family of transcription factors, NFκB regulates hundreds of genes, but these are only a fraction of the genes whose expression is influenced by sustained PKC activation. This becomes evident from the large number of differentially expressed genes and altered signaling pathways after 3 h of PMA treatment in the transcriptomic analysis (Fig. 5). Our sequencing data support key findings that were made using pharmacological inhibitors: 1) PMA upregulates the NFκB pathway and the MAPK pathway. 2) TPCA1+NIKSMI1 were very effective in inhibiting the NFκB pathway. In fact, they almost restored it to control level, while hardly influencing any other pathway. Notably, EGFR and MAPK, which both include ERK1/2^116^, were still highly enhanced in the presence of these inhibitors. This provides strong evidence that the structural and functional effects observed with TPCA1+NIKSMI1 were a result of gene expression regulated by NFκB, even though MKK1 inhibitors were also effective (see 4.3). Furthermore, we performed a GSEA to elucidate pathways and cellular functions not included in the PROGENy database. We added a set of t-tubule-associated genes which was derived from a recent study investigating the t-tubule proteome^51^. It turned out that gene sets related to cardiac disease, endocytosis, integrins and the cytoskeleton were significantly altered by PMA and reversed by blocking NFκB. Importantly, NFκB inhibition yielded a highly significant change in the t-system-associated genes, which fits well to the stabilizing effect on the t-tubules. Surprisingly, however, the result showed an overall downregulation of these genes. T-system-related proteins may be lost during t-tubule degradation, which could create signals for increased expression in the sense of compensatory mechanisms. One should also keep in mind that most likely only few of the approximately 150 t-system-associated genes are responsible for t-tubule development, maintenance and degradation.

In sum, these results confirm that t-system-related genes are affected and suggest that processes related to the cytoskeleton and to endocytosis are involved in t-tubule loss, of which the former fits well to published studies^17,105^, whereas the latter may open a new perspective on t-tubule remodeling and provide the link between the cytoskeleton and the t-system, as endocytosis-related processes heavily rely on the cytoskeleton.

To narrow down the genes that may be of particular relevance for t-system remodeling, we created intersections of the list of t-system-associated genes and the genes oppositely regulated by PMA and PMA + NFκB inhibition (Fig. 5G). The identified genes include dysferlin (DYSF), which has been demonstrated to be involved in membrane repair and t-tubule proliferation and organization^117^. Interestingly, other studies showed that dysferlin interacts with annexins, BIN1 and the actin cytoskeleton during membrane repair^118^ and has been related to integrin-induced macropinocytosis in neonatal rat cardiomyocytes^119^, which fits well to the differential regulation of ITGB1, as it has been reported to supress autophagy, induce macropinocytosis and control cardiomyocyte behaviours^120,121^. However, dysferlin was upregulated by PMA, as well as BIN1, suggesting a compensatory role rather than a primary cause of t-tubule loss. FHL2 (Four And A Half LIM Domains 2) was the only t-system-related gene that was downregulated by PMA and upregulated again after NFκB inhibition. FHL2 is ubiquitously expressed, belongs to the genes with highest expression in the adult human ventricular myocytes^122^ and is involved mainly in protein-protein interactions^123^. Clinically, FHL2 is relevant for cancer^124^, plays a role in cardiac hypertrophy and mutations have been identified in patients with familial dilated cardiomyopathy^125^. Based on these findings, we suggest to explore the role of FHL2 also in t-system remodeling.

### 4.5. Effects on Excitation-Contraction Coupling

As expected, the severe loss of t-tubules affected the integrity of cardiac dyads, as visible from increased RyR-LTCC distances, which resulted in inefficient, slowed and delayed intracellular Ca^2+^ release and decreased contractility (Fig. 6, Fig. **S6**). Blocking NFκB rescued cardiomyocyte function (Ca^2+^ signaling and contractility), which underscores the importance of t-tubules for excitation-contraction coupling and provides evidence for consequences of NFκB activation on adult cardiomyocyte function. Interestingly, the combination of PMA and TPCA1+NIKSMI1 resulted in some parameters being even better than in the control group (Fig. 6D), suggesting that NFκB is activated spontaneously in cell culture and may regulate additional EC coupling features.

Surprisingly, JPH2 density was not recovered by blocking NFκB, suggesting that JPH2 is not directly involved in t-system maintenance. This fits to our previous findings, where glucocorticoids had no effect on JPH2 despite stabilizing the t-system against spontaneous degradation in cell culture^15^ and to studies suggesting the main role of JPH2 in modulating RyR function and RyR-LTCC interaction rather than directly stabilizing the t-tubule membrane^126^.

BIN1, a protein known to orchestrate t-tubule formation in developing cardiomyocytes^123^ and to be associated with endocytic processes^127^, showed disrupted organization and lower protein levels upon PMA treatment. This effect is revoked when NFκB inhibitors are added and t-tubules are maintained. However, our transcriptomic analysis revealed increased BIN1 expression in the PMA group, while the addition of TPCA1+NIKSMI1 had no effect. We therefore propose that increased BIN1 mRNA expression may constitute a compensatory reaction when t-tubules are lost rather than a cause of PMA-induced t-system degradation.

### 4.6. Macropinocytosis as the Mechanism of T-Tubule Internalization

Utilizing fixable dextran and a fixable lipophilic membrane dye, we observed that t!zltubules are internalized into the cytosol of cardiac myocytes, leading to their disappearance (Fig. 7). Importantly, we showed that the rate of internalization correlates linearly with the rate of t-tubule loss, i.e. all compounds that prevented t-system deterioration also reduced the amount of internalized dextran. By varying the timing of membrane dye application, we demonstrated that the dextran-labelled structures are enclosed by membranes that were originally contiguous with the plasma membrane but are no longer connected (Fig. S7). Internalization of the plasma membrane, including its components and extracellular molecules can be caused by different types of endocytosis. These can be divided into their dependence on different proteins, such as clathrin, dynamin, caveolin or actin. Because of the large size of the internalized structures (up to several micrometers), we suspected a macropinocytosis-related process. Macropinocytosis is an actin-dependent, clathrin-and dynamin-independent large-scale endocytic process that internalizes large volumes of extracellular fluid^128^. The formation of macropinosomes is mediated by membrane ruffling and closure, which requires class I PI3K-dependent actin polymerization ^95–97^ and may also require myosin I and non-muscle myosin II. We tested several known inhibitors of macropinocytosis, including PI3K class I inhibitors, actin inhibitors and a myosin I inhibitor, which all were effective in reducing internalized t-tubule membranes. Dynasore, an inhibitor of dynamin, however, had little effect (Figs. 8 and S8). PI-103 and wortmannin, which are highly specific and potent inhibitors of class I PI3Ks^74^, prevented the dextran uptake. This strongly suggests that t-tubules in isolated cardiomyocytes are lost by a macropinocytic process. It also fits to reports that PMA upregulates macropinocytosis in fibroblasts^129^ and neutrophils^130^, suggesting that PKC-NFkB signaling could drive membrane remodeling not only in cardiomyocytes.

The effectiveness of PI3K inhibitors leads to the question whether the PI3K-PDK-AKT signaling pathway is involved, especially when inspecting Fig. 5D, which shows that in addition to NFκB also the PI3K pathway is upregulated by PMA and brought back to control levels by the addition of NFκB inhibitors. However, there are two lines of evidence suggesting that PI3K-AKT is not involved. First, the inhibition of AKT by AKTI1/2 or of PDK by GSK2334470 (Fig. S5G) did not prevent t-tubule loss. Second, PKD inhibitors were very effective in rescuing the t-system, although PKD is not known to activate AKT. Therefore, it is more likely that the inhibition of PI3K class I prevented macropinosome formation directly. The finding that t-tubules are internalized fits to previous work describing that, during long-term cardiomyocyte culture, t-tubules are slowly internalized by sequential inward pinching^131^. It would even fit to the observation that some t-tubules fail to conduct action potentials after the application of voltage-sensitive membrane stainings^132^, as it seems possible that some t-tubules may have been internalized and lost contact to the surface membrane.

An important question is what fate t-tubules adopt once internalized into the cytosol. As shown in Fig. 8, the dextran content decreases following washout in an exponential manner. This can be explained by several possibilities: First, the internalized t-tubule vesicles might reintegrate into the plasma membrane, leading to the release of dextran into the extracellular medium. Alternatively, the vesicles could be degraded intracellularly, which would result in the dextran being dispersed into the cytosol, where it may no longer be detectable as a signal. Additionally, chemical degradation of the dextran or fluoropohre could occur, leading to a loss of fluorescence. Spontaneous chemical degradation, however, would not explain different decay rates between PMA and CTRL. Notably, PMA-treated cells exhibit a more rapid decay compared to controls, yet they maintain a higher steady-state level and a greater maximum uptake velocity, suggesting an elevated turnover rate. Although we cannot exclude the possibility that some t-tubules are re-integrated into the plasma membrane, the strong correlation between dextran uptake and t-tubule loss implies that at least a portion of the internalized t-tubules is degraded, possibly via lysosomal fusion. Further research is necessary to elucidate the processes occurring after t-tubule internalization.

### 4.7. Clinical and Pathophysiological Relevance

PKC plays diverse roles in the heart ^133^ and its expression varies depending on disease stage and pathology ^134^. However, our findings align well with the concept of chronic PKC activation in heart failure^135^, particularly in relation to t-system remodeling. Our data suggest that chronic activation of PKC and NFκB in the heart drives cytoskeletal changes and t-tubule disorganization as a key consequence. Since adverse t-system remodeling is a hallmark of heart failure progression and a potential therapeutic target, this mechanism is of clinical importance. Notably, hearts with a disrupted t-system show poor prognosis^7,103^. A therapeutic goal is to prevent or even reverse t-tubule disorganization to preserve contractile function in failing myocardium. Our study suggests the contemplation of two potential approaches: targeting PKC–NFκB activation or/and processes related to micropinocytosis and membrane-traffic in cardiac myocytes. Chronic cardiac NFκB activation is a sign of inflammation, drives cardiac remodeling, such as hypertrophy and fibrosis and is associated with poor prognosis^136^. Thus, cardiac or cardiomyocyte-specific inhibition of this pathway may offer therapeutic benefits^137^ which also comprise the reduction of t-system remodeling. Secondly, we demonstrate that blocking macropinocytosis in isolated cardiomyocytes reduces the internalization and loss of t-tubules. To explore translational potential, we tested FDA-approved macropinocytosis inhibitors^100^ (Fig. S8), which showed some effect but only modest efficacy. Nevertheless, we suggest further research to identify more effective drugs that can halt t-system degradation.

### 4.8. Limitations and Future Directions

Our study provides novel insights into the PKC–PKD–NFκB pathway and mechanisms of t-system remodeling. However, some limitations should be acknowledged. Our experiments relied predominantly on isolated cardiomyocytes. It is well known that cell isolation procedures and cell culture can induce t-tubule loss. Thus, the stability of t-tubules in isolated cardiomyocytes is already affected under baseline conditions, although we showed that this spontaneous degradation can be blocked by NFκB inhibition (Fig. S4). To overcome this limitation, we also used rabbit and rat myocardial slice culture, in which t-tubules were shown to remain stable^138^. Furthermore, we observed that the extent and rate of t-system breakdown differ among rat, rabbit, and human cardiomyocytes. In tissue cultures, TT loss occurs more gradually, likely reflecting higher t-tubule stability when anchored to the extracellular matrix. In addition, while PMA-induced PKC activation is a robust model for inducing t-system degradation, it represents an overactivation of PKC. Receptor-mediated agonists also increased the internalization rate, albeit to a lesser extent. This suggests they hypothesis that PKC induces t-tubule loss also in vivo occurs, but not as quickly as shown here. Testing hypothesis would require experiments beyond the scope of this study. Transcriptomic analysis identified several candidate genes that may be involved in this process, although one needs to acknowledge that the gene sets used might not be fully comprehensive. Lastly, we acknowledge the inherent limitations of pharmacological inhibitors, including off-target effects. To mitigate this, however, we used inhibitors validated in large kinase panels^74–76^ where possible, employed multiple inhibitors per target, and corroborated key findings with RNA sequencing data.

In summary, despite the inherent limitations of in-vitro models, this study uncovers a novel PKC–PKD–NFκB–driven, actin-dependent macropinocytic mechanism underlying t-tubule loss in cardiomyocytes. These findings deepen our understanding of the molecular basis of heart failure and suggest broader relevance beyond the heart, as the signaling pathways and mechanisms identified here may also operate in non-muscle and non-cardiac cells. Membrane internalization and degradation are implicated in diverse pathologies, including neurodegeneration, immune cell dysfunction, and cancer, where similar actin- and kinase-dependent processes contribute to cellular remodeling and disease progression. Both NFκB signaling and macropinocytosis can promote survival and metabolic adaptation of tumor cells^139,140^. Thus, future studies could explore whether PKC–PKD–NFκB–driven membrane remodeling mechanisms, as identified here in cardiomyocytes, also operate in non-cardiac systems and cancer cells. Future research should also dissect the isoform-specific contributions of PKC and PKD to membrane remodeling, explore the crosstalk between ERK1/2 and NFκB pathways and further elucidate the molecular details of t-tubule endocytosis and degradation.

## Acknowledgements

We would like to thank to thank Dr. Hendrik Milting from the Heart and Diabetes Center NRW, Erich and Hanna Klessmann Institute, University Hospital of the Ruhr-University Bochum, Bad Oeynhausen, Germany, for his help with the acquisition of human heart samples. We would like to thank C. Grüninger and L. McCargo for their highly valuable technical support. A-K.M.P, J.W., L.K.K. and P.A. performed this work in (partial) fulfilment of the requirements for obtaining the degree “Dr. med.”.

---

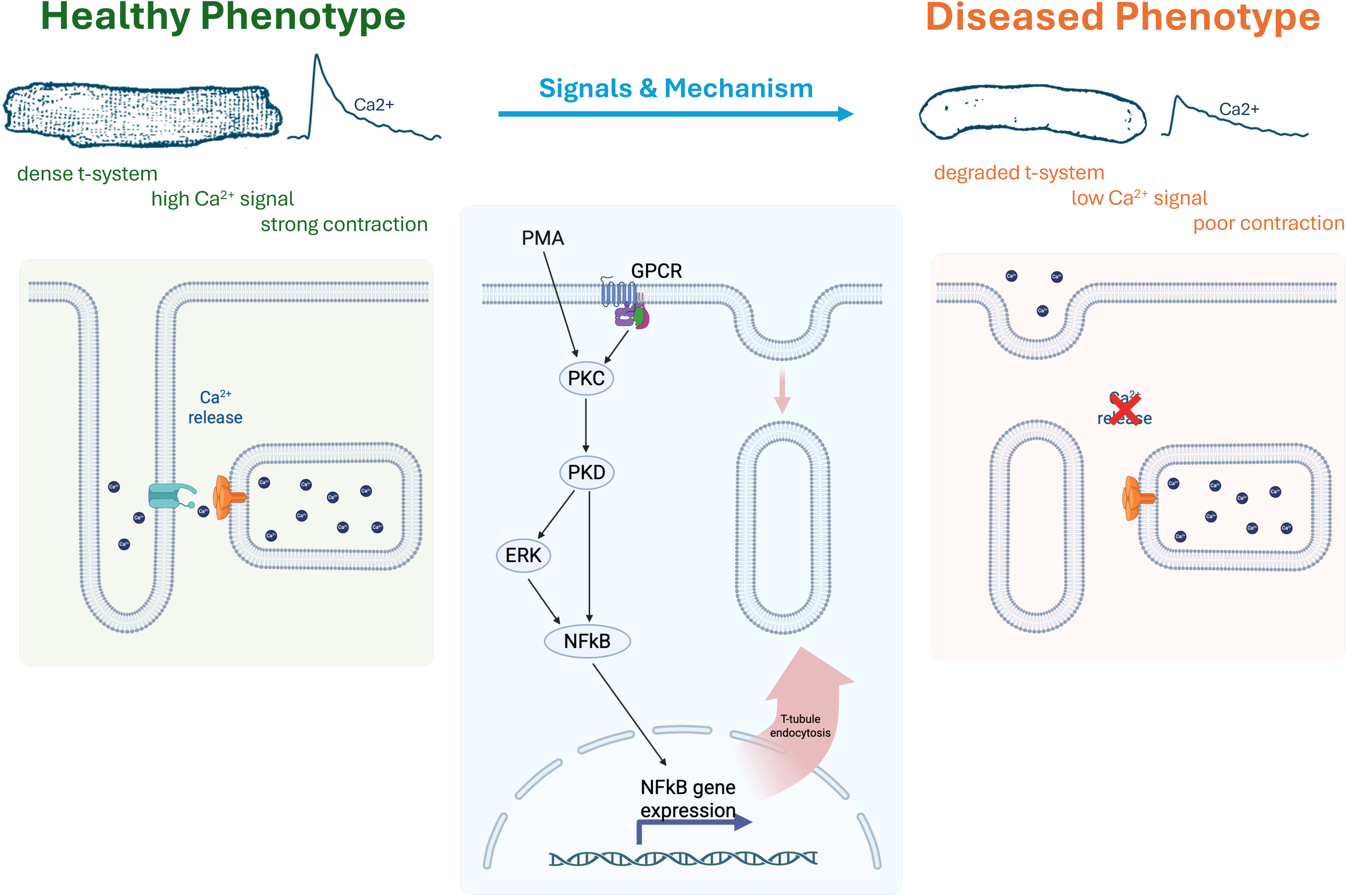

