## Supplemental Figures for "PKC Promotes T-Tubule Membrane Loss by Activating a PKD– NFκB Endocytic Pathway"

### Figure S1

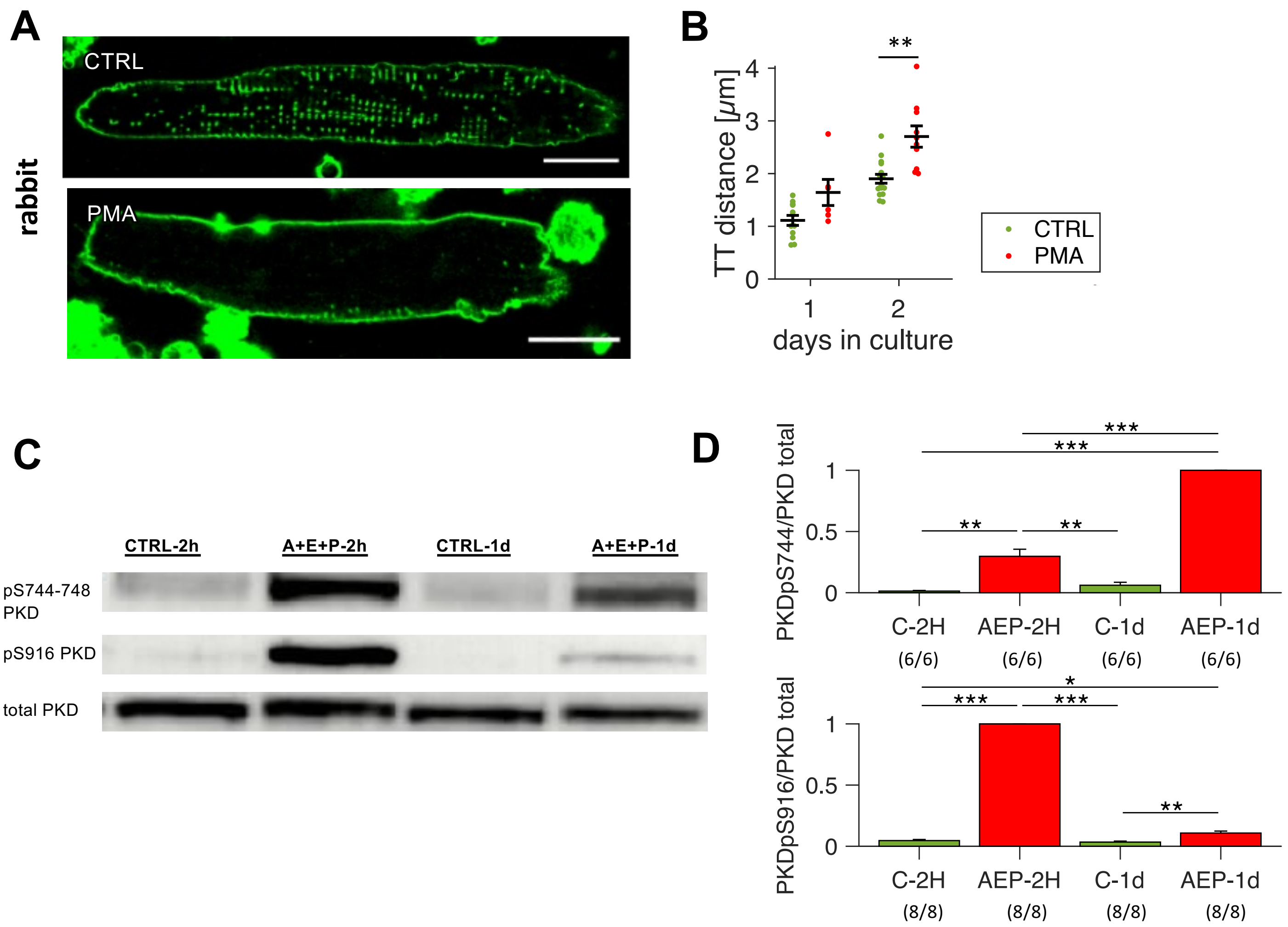


**Fig. S1. (A)** Confocal microscopic images of Di8-ANEPPS-stained example isolated rabbit cardiomyocytes cultured for one day with either vehicle (CTRL) or 50 nM PMA. Scale bar: 20 µm. **(B)** Mean intracellular t-tubule (TT) distance in PMA and CTRL rabbit cells after one and two days in culture. Number of cells / hearts: 12/2 and 16/2 in CTRL, 6/1 and 10/1 in PMA after 1d or 2d in culture, respectively, ** $p<0.05$; Welch's t-test with multiple comparison correction. **(C)** Western Blots of rat cardiomyocytes treated for two hours (2h) or one day (1d) with vehicle (CTRL) or 1µM angiotensin II + 100nM endothelin 1 + 10µM phenylephrine (AEP). Bands show total protein kinase D (PKD) and the PKC-specific phosphorylation site of PKD at S744-748 and the PKD autophosphorylation site at S916. **(D)** Quantification and statistical analysis of the Western blots. Band intensities were normalized to total PKD, Ponceau staining intensity and AEP on each blot. * $p<0.05$; Welch's t-test with multiple-comparison correction.

### Figure S2

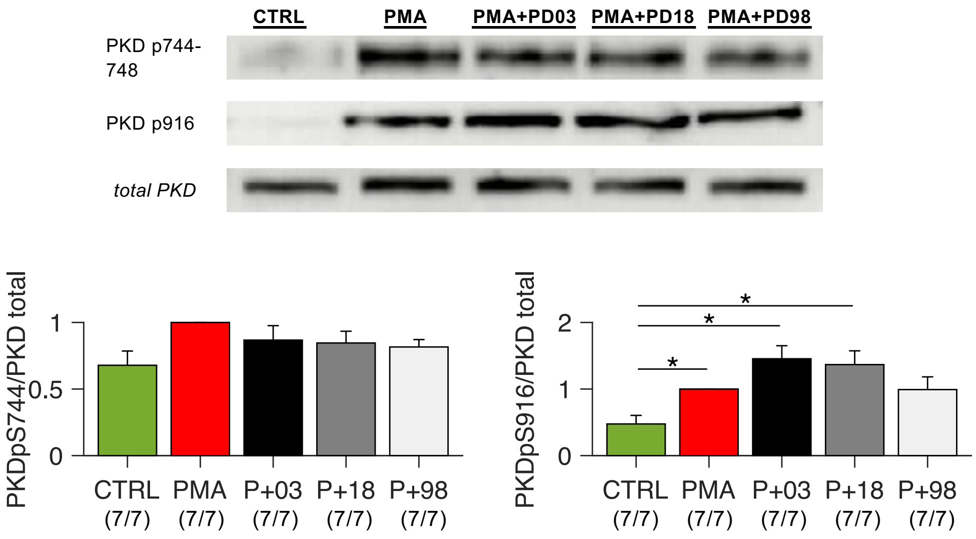


**Fig. S2: PKC/PKD activation by MKK1**

Effect of inhibitors for MKK1 PD0325901 (100 nM), PD184352 (1 µM) and PD98059 (1 µM) on PKC and PKD activation. The compounds were incubated for 1h before PMA addition and further cultured for 1 day before freezing. Phosphorylation of PKD at the PKC-specific (pS744-748) and autophosphorylation (pS916) site were assessed by Western blotting. Band intensities were normalized to total PKD levels and to Ponceau staining intensity as loadin control. The results were normalized to the respective normalized PMA band intensities. Numbers below the graphs indicate the number of technical / biological replicates (evaluated blots / heart cell isolations). * $p < 0.05$, unpaired, non-equal variance t-test, cells from matched cell isolations.

Figure S3:

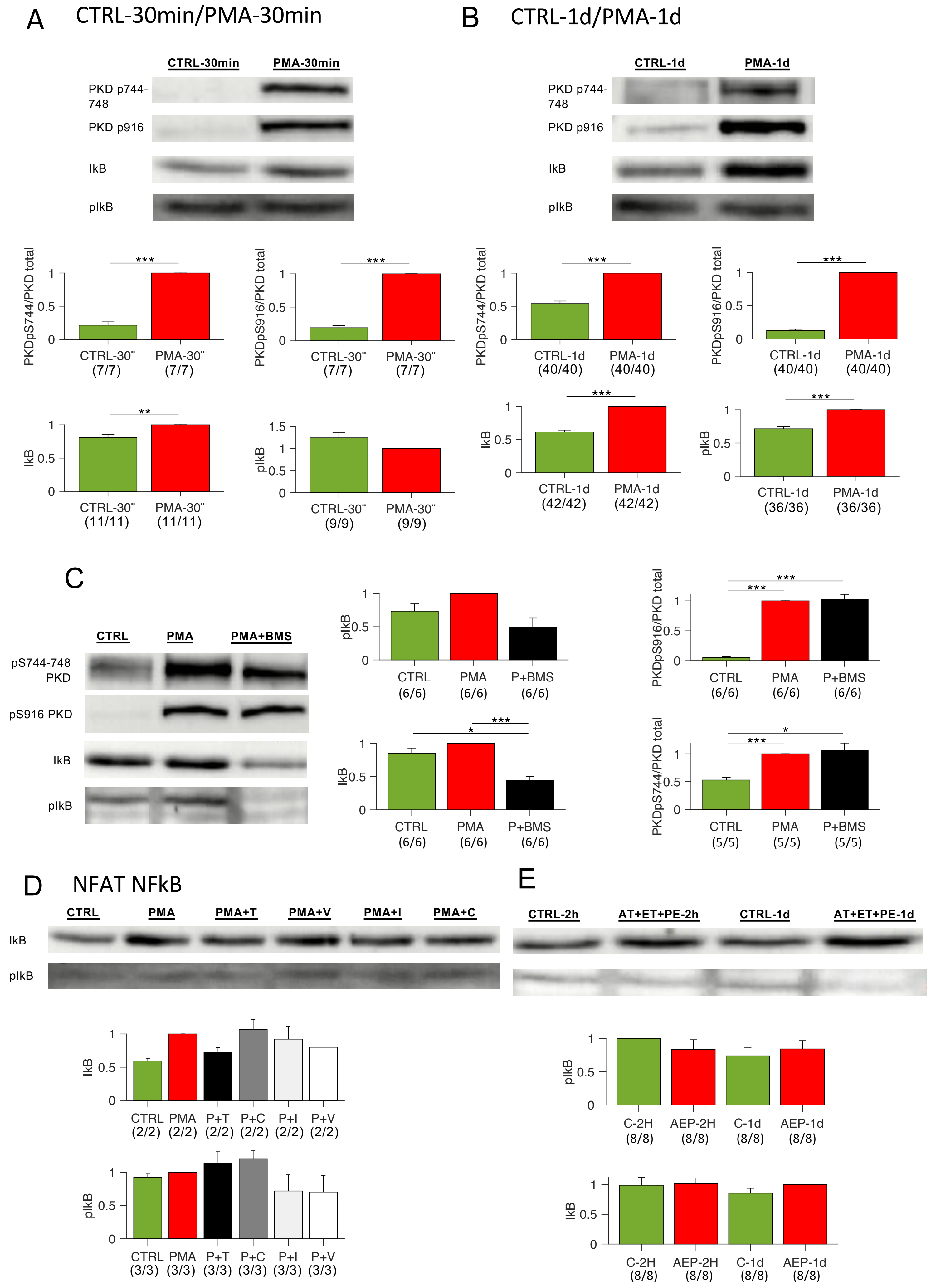
A
CTRL-30min/PMA-30min
CTRL-30min
PMA-30min
PKD p744-748
PKD p916
IkB
pIkB
PKDpS744/PKD total
PKDpS916/PKD total
IkB
pIkB
CTRL-30'' (7/7)
PMA-30'' (7/7)
CTRL-30'' (11/11)
PMA-30'' (11/11)
CTRL-30'' (9/9)
PMA-30'' (9/9)
B
CTRL-1d/PMA-1d
CTRL-1d
PMA-1d
PKD p744-748
PKD p916
IkB
pIkB
CTRL-1d (40/40)
PMA-1d (40/40)
CTRL-1d (42/42)
PMA-1d (42/42)
CTRL-1d (36/36)
PMA-1d (36/36)
C
CTRL
PMA
PMA+BMS
pS744-748 PKD
pS916 PKD
IkB
pIkB
CTRL (6/6)
PMA (6/6)
P+BMS (6/6)
CTRL (5/5)
PMA (5/5)
P+BMS (5/5)
D
NFAT NFkB
CTRL
PMA
PMA+T
PMA+V
PMA+I
PMA+C
CTRL (2/2) PMA (2/2) P+T (2/2) P+C (2/2) P+I (2/2) P+V (2/2)
CTRL (3/3) PMA (3/3) P+T (3/3) P+C (3/3) P+I (3/3) P+V (3/3)
E
CTRL-2h
AT+ET+PE-2h
CTRL-1d
AT+ET+PE-1d
C-2H (8/8)
AEP-2H (8/8)
C-1d (8/8)
AEP-1d (8/8)

**Fig. S3:**

Effects of PMA on PKC, PKD and NFkB activation after 30 min **(A)** and 1 day **(B)**; phosphorylation of PKD at the PKC-specific (pS744-748) and autophosphorylation (pS916) site, as well as total and phosphorylated IkB levels were assessed by Western Blot. **(C)** Effects of broad-spectrum NfkB inhibitor BMS-34554 20µM on PMA-induced activation of PKC, PKD and NFkB; phosphorylation of PKD at the PKC-specific (pS744-748) and autophosphorylation (pS916) site, as well as total and phosphorylated IkB levels were assessed by Western Blot. **(D)** Effect of NFAT inhibitors tacrolimus 5µM, VIVIT 10µM, INCA 5µM or ciclosporin A 1µM on NFkB activation, total and phosphorylated IkB levels were assessed by Western Blot. **(E)** Effect of G-coupled receptor agonists 1µM angiotensin II, 100 nM endothelin 1 and 10 µM phenylephrine on NFkB activation, total and phosphorylated IkB levels were assessed by Western Blot.

Statistical test: paired t-test. P-values after correction for multiple comparisons: *$p < 0.05$; **$p < 0.01$; ***$p < 0.001$.

### Supplemental Figure S4:

A

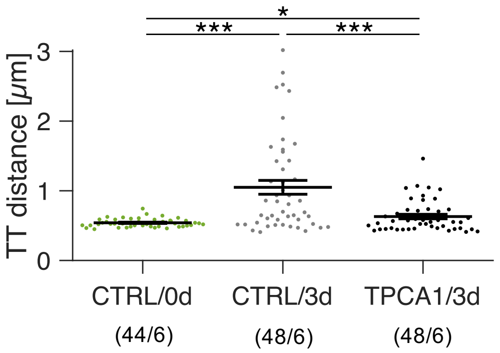


**Supplemental Fig. S4: Effect of NFkB inhibition on spontaneous t-tubule loss in cell cutlure**
Rat cardiac myocytes were stained with Di8 ANEPPS and imaged by confocal microscopy TPCA1 either immediately after cell isolation (CTRL/0d) or after three days in culture treated with vehicle (CTRL/3d) or with the NFkB inhibitor TPCA1 (TPCA1/3d). Intracellular t-tubule (TT) distances was used a measure of t-system degradation. * p<0.05, *** p<0.001 (unpaired Welch's t-test, multiple comparison correction). Numbers below group names indicate cells/animals.

### Figure S5

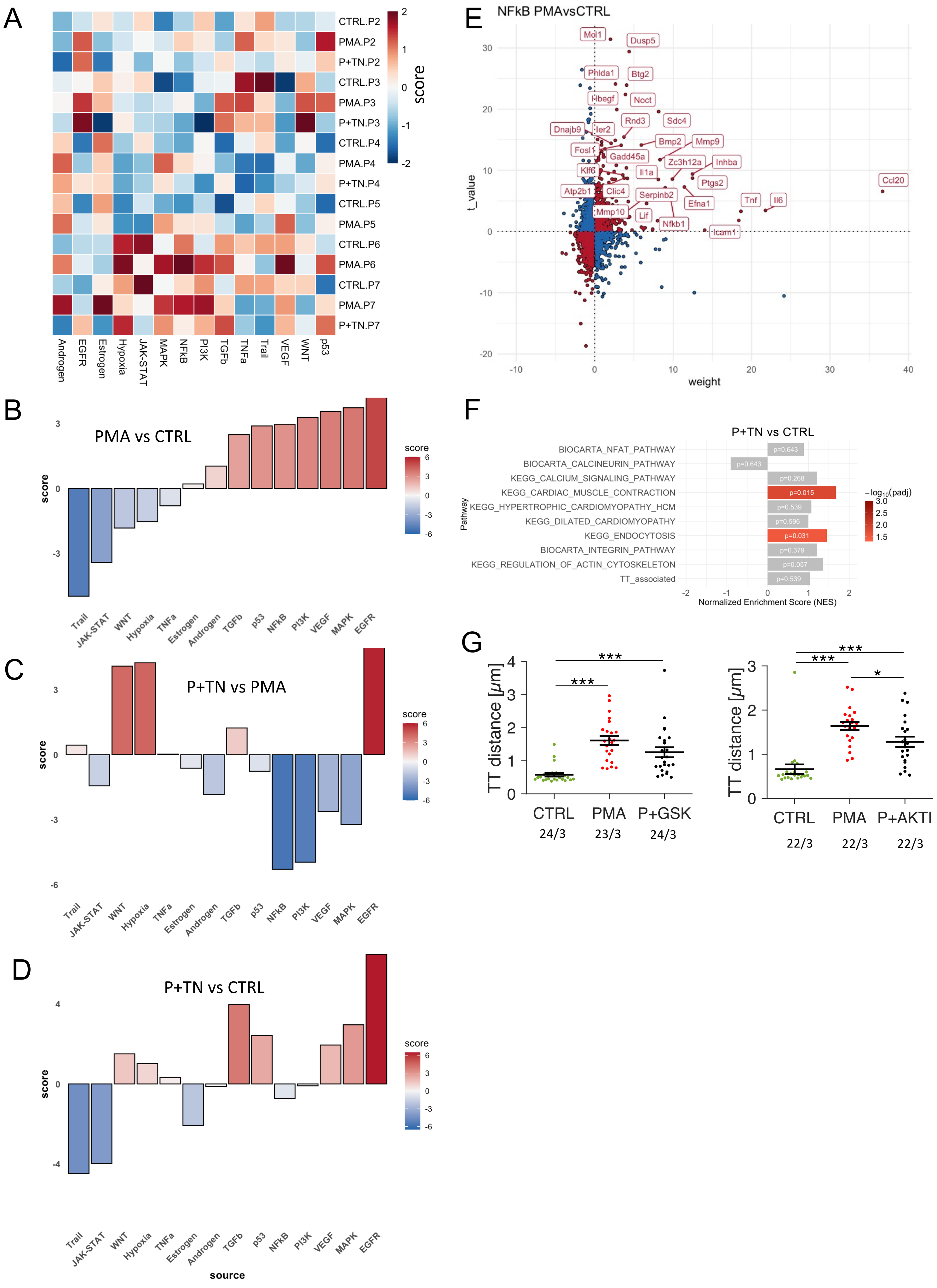

A
CTRL.P2
PMA.P2
P+TN.P2
CTRL.P3
PMA.P3
P+TN.P3
CTRL.P4
PMA.P4
P+TN.P4
CTRL.P5
PMA.P5
CTRL.P6
PMA.P6
CTRL.P7
PMA.P7
P+TN.P7
score
Androgen
EGFR
Estrogen
Hypoxia
JAK-STAT
MAPK
NFkB
PI3K
TGFb
TNFa
Trail
VEGF
WNT
p53
B
PMA vs CTRL
C
P+TN vs PMA
D
P+TN vs CTRL
score
source
Trail
JAK-STAT
WNT
Hypoxia
TNFa
Estrogen
Androgen
TGFb
p53
NFkB
PI3K
VEGF
MAPK
EGFR
E
NFkB PMAvsCTRL
t_value
weight
Mol1
Dusp5
Phlda1
Btg2
Hbegf
Noct
Rnd3
Sdc4
Dnajb9
Ier2
Bmp2
Mmp9
Fosl1
Gadd45a
Zc3h12a
Inhba
Klf6
Il1a
Ptgs2
Ccl20
Atp2b1
Clic4
Serpinb2
Efna1
Tnf
Il6
Mmp10
Lif
Nfkb1
Icam1
F
P+TN vs CTRL
Pathway
BIOCARTA_NFAT_PATHWAY
BIOCARTA_CALCINEURIN_PATHWAY
KEGG_CALCIUM_SIGNALING_PATHWAY
KEGG_CARDIAC_MUSCLE_CONTRACTION
KEGG_HYPERTROPHIC_CARDIOMYOPATHY_HCM
KEGG_DILATED_CARDIOMYOPATHY
KEGG_ENDOCYTOSIS
BIOCARTA_INTEGRIN_PATHWAY
KEGG_REGULATION_OF_ACTIN_CYTOSKELETON
TT_associated
p=0.643
p=0.643
p=0.268
p=0.015
p=0.539
p=0.596
p=0.031
p=0.379
p=0.057
p=0.539
Normalized Enrichment Score (NES)
−log10(padj)
G
TT distance [μm]
CTRL
PMA
P+GSK
24/3
23/3
24/3
CTRL
PMA
P+AKTI
22/3
22/3
22/3

**Fig. S5**

**(A)** Heatmap showing the score of a PROGENy pathway analysis in individual CTRL, PMA and PMA+NIKSMI1+TPCA1 (P+TN) samples, using the log-normalized counts. The lines of the tree on the left indicate the Euclidean distance between the samples. The score was obtained by column-wise scaling (subtraction of mean, followed by division by the standard deviation). P1-P7 indicate the batches (hearts). **(B-D)** PROGENy pathway analysis, using the Wald statistics value of the differential expression analysis of PMA vs CTRL, P+TN vs PMA and P+TN vs CTRL. **(E)** Genes with their weights from the PROGENy NFkB pathway model and their *t*-value from the multilinear model. Genes contributing to the downregulation of the pathway are shown as blue circles, genes contributing to upregulation as red circles. The 30 genes with the highest score (absolute value of weight multiplied with *t*-value) are labelled, i.e. the genes contributing most to upregulation of the NFkB pathway in PMAvsCTRL. **(F)** Gene set enrichment analysis (GSEA) of P+TN vs CTRL. The direction and magnitude indicate the normalized enrichment score, while the color indicates the adjusted p value. **(G)** Mean t-tubular (TT) distance in rat cardiomyocytes treated with vehicle (CTRL), 50 nM PMA or 50 nM PMA + PDK inhibitor GSK2334470 (P+GSK) or PMA + AKT (PKB) inhibitor AKTI-1/2 (P+AKTI). * $p<0.05$, *** $p<0.001$; unpaired, non-equal variance (Welch's) t-test; cells from matched cell isolations

### Figure S6

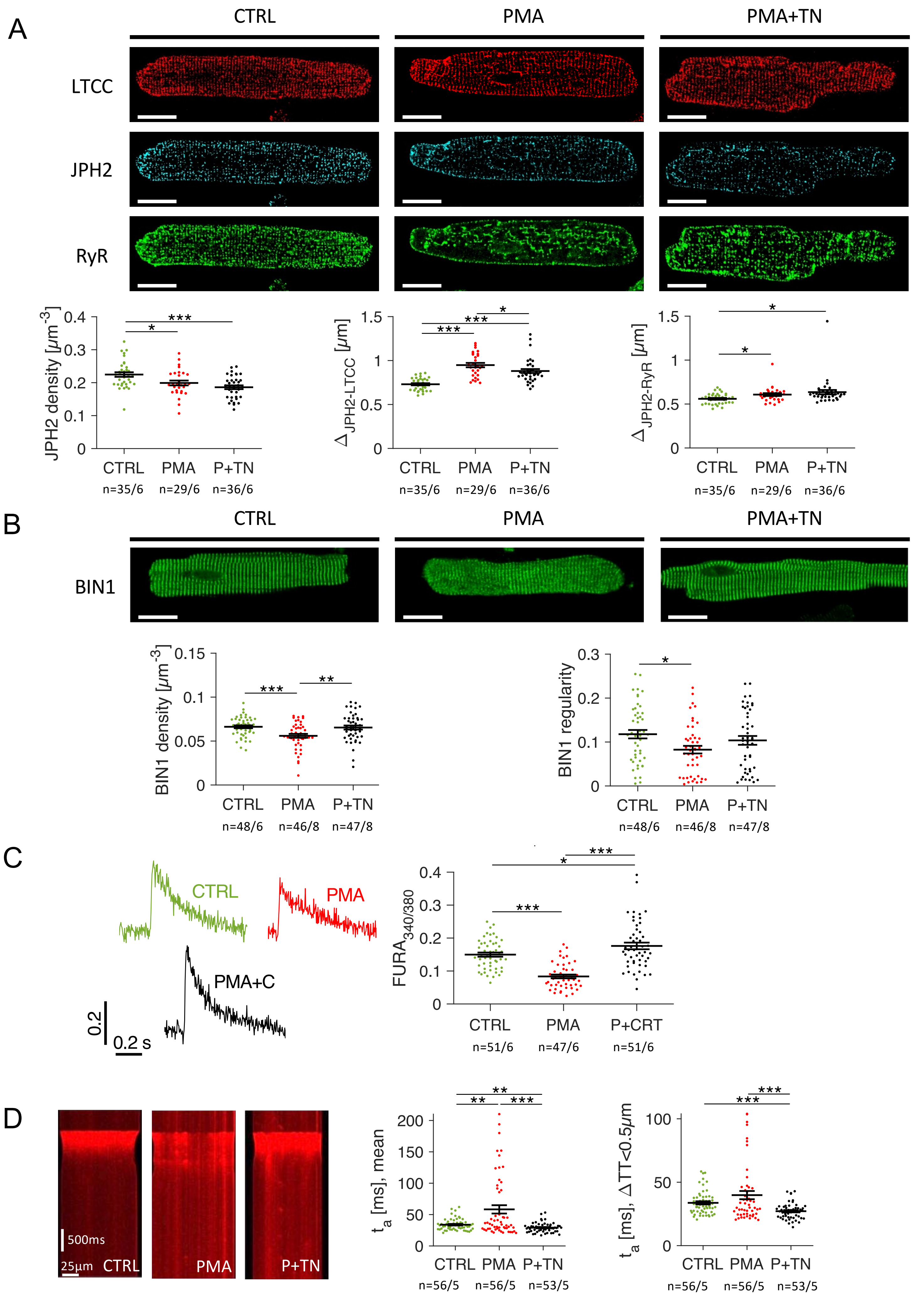

A
CTRL
PMA
PMA+TN
LTCC
JPH2
RyR
JPH2 density [μm⁻³]
0.4
0.3
0.2
0.1
0
***
*
CTRL
PMA
P+TN
n=35/6
n=29/6
n=36/6
Δ JPH2-LTCC [μm]
1.5
1
0.5
0
***
***
*
Δ JPH2-RyR [μm]
B
BIN1
BIN1 density [μm⁻³]
0.15
0.1
0.05
***
**
n=48/6
n=46/8
n=47/8
BIN1 regularity
0.3
0.2
0.1
C
CTRL
PMA
PMA+C
0.2
0.2 s
FURA340/380
P+CRT
n=51/6
n=47/6
D
500ms
25μm
ta [ms], mean
200
150
100
50
n=56/5
n=56/5
n=53/5
ta [ms], ΔTT<0.5μm

**Fig. S6**

**(A)** Immunofluorescent images of rat cardiomyocytes treated for 24 h with vehicle (CTRL), PMA or 50 nM PMA + TPCA1 + NIKSMI1 (P+TN). Cells were stained for L-type $Ca^{2+}$ channels (LTCC, red), junctophilin 2 (JPH2, cyan) and ryanodine receptors (RyR, green). The graphs below the images show the quantification of JPH2 cluster density (voxels positive for JPH2 divided by cell volume), the mean centroid distance of JPH2 clusters to their closest LTCC cluster (ΔJPH2-LTCC) and to their closest RyR cluster (ΔJPH2-RyR). **(B)** Confocal images of rat cells stained for BIN1 and the quantification of BIN1 volume density as well as BIN1 regularity, calculated as the spectral density, using the Fourier transform of the raw signal. **(C)** Example traces of the $Ca^{2+}$ signal of Fura-2-loaded cells at 1 Hz pacing rate at 35 ºC. Cells were treated for 24 h with vehicle (CTRL), PMA or PMA + the PKD inhibitor CRT (P+CRT0066101). Mean amplitude of the $Ca^{2+}$ transients are shown next to the traces. **(D)** Confocal line scans of rat cardiomyocytes cultured for 24 h under CTRL, PMA or P+TN conditions, loaded with FLUO4 as $Ca^{2+}$ indicator. Mean activation time (ta), identified as the time of maximum $Ca^{2+}$ signal upstroke velocity of the whole line and restricted to the voxels within the lines that were very close to the closest t-tubule (TT distance < 0.5µm).

* $p<0.05$, ** $p<0.01$, *** $p<0.001$, unpaired non-equal variance (Welch's) t-test, n/N indicates number of cells or measurements / number of biological replicates (hearts). Cells from matched cell isolations.

### Figure S7

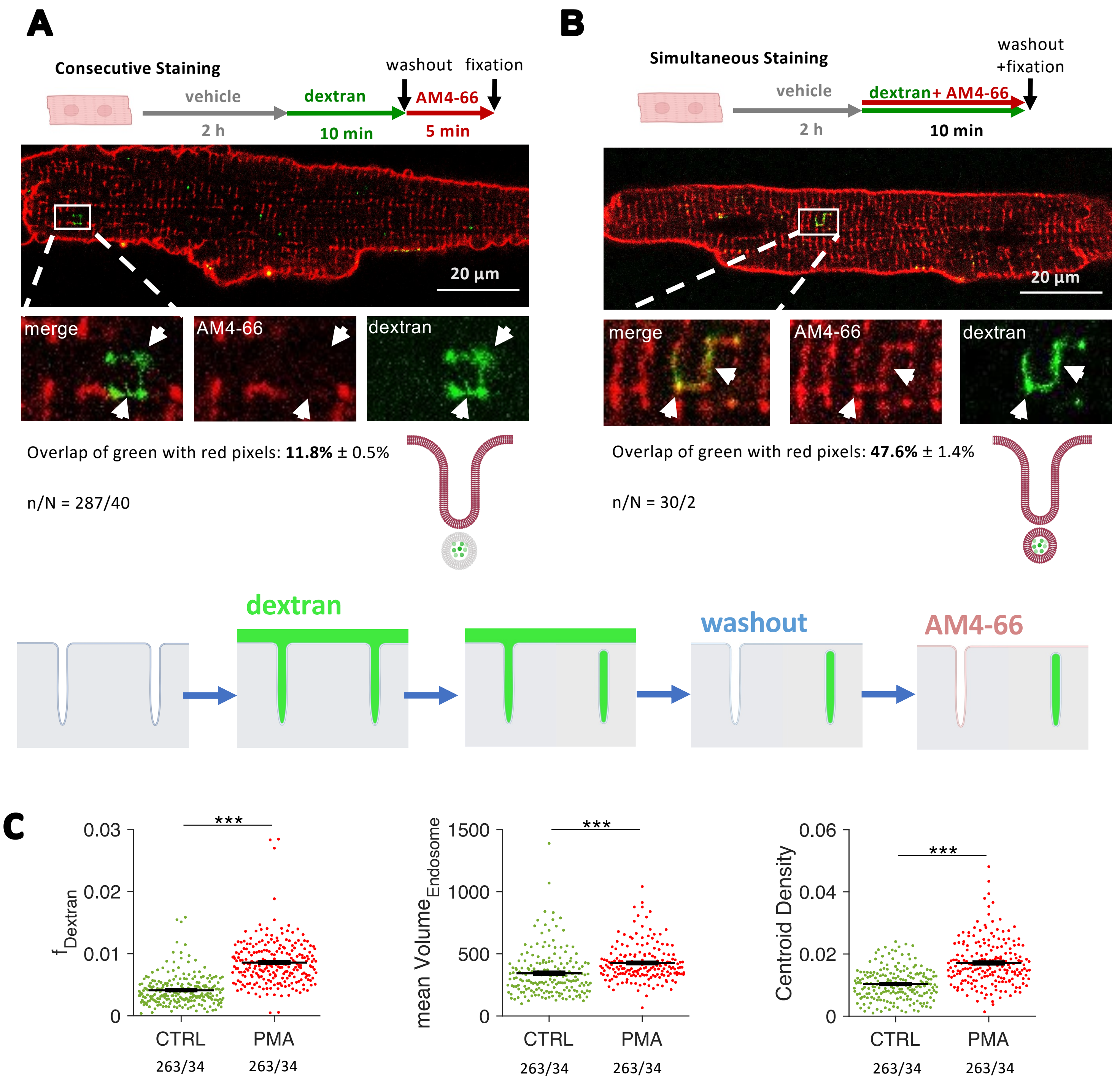


**Fig. S7: Mechanisms of t-tubule internalization**
**(A + B)** Methodological approach and confocal images displaying an overview, magnified view of channels stained with the membrane stainin AM4-66 (red) and 10 kDa dextran (green) and graphical representation of dextran internalization and staining. **(A)** Result after consecutive staining. **(B)** Result after simultaneous staining, used to analyze the overlap between AM4-66 and dextran staining. The overlap of pixels positive for both stains, divided by dextran-positive pixels, was quantified (mean ± standard error). **(C)** Analysis and quantification of parameters related to internalization in cells incubated with vehicle (CTRL) and PMA (50 nM) for 2 hours.
Statistical test: paired t-test. ***p < 0.001.

Figure S8

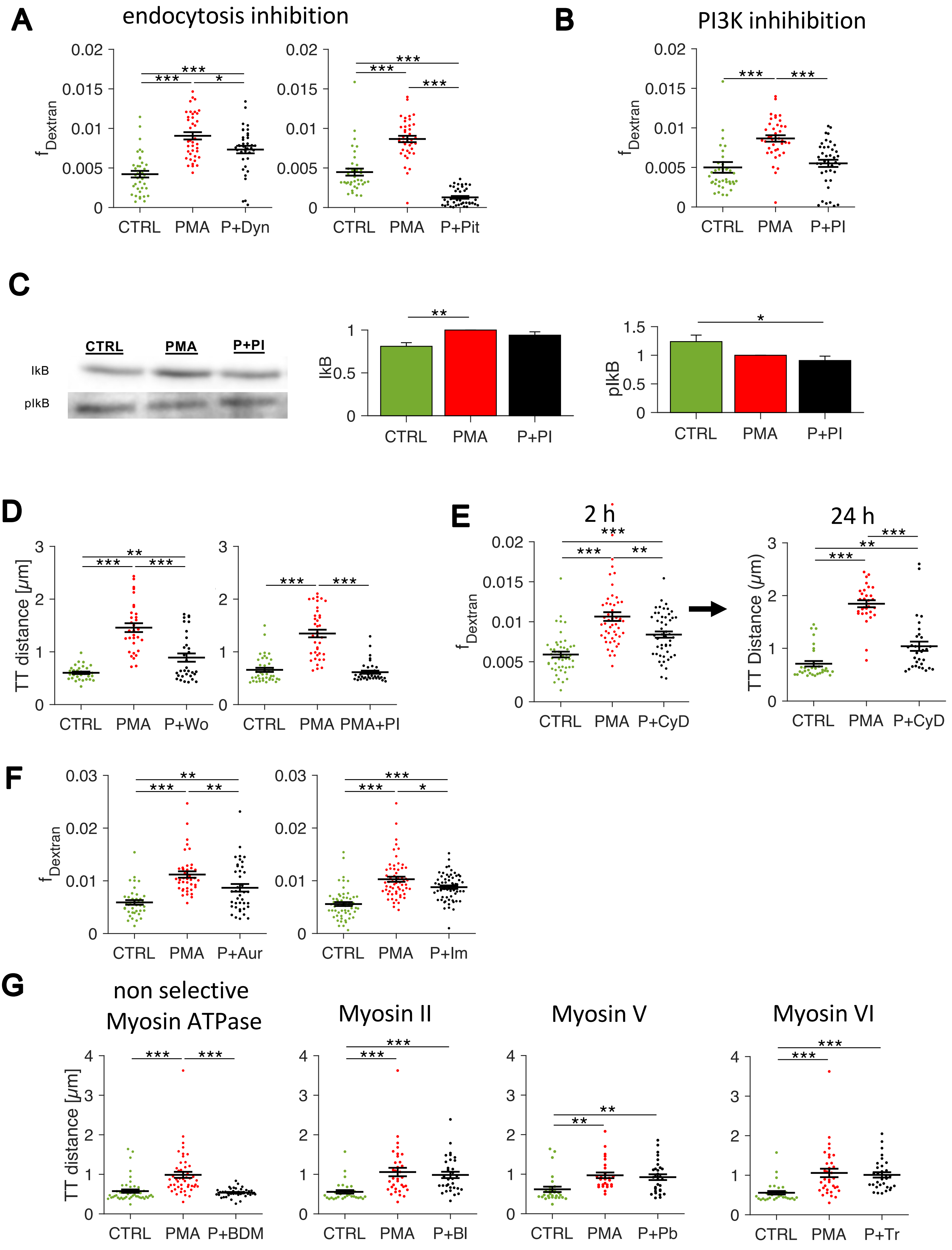

A
endocytosis inhibition
B
PI3K inhihibition
C
CTRL
PMA
P+PI
IkB
pIkB
D
E
2 h
24 h
F
G
non selective Myosin ATPase
Myosin II
Myosin V
Myosin VI

**Fig. S8: T-tubule internalization via macropinocytosis**

**(A)** Quantification of the volume ratio of internalized dextran in cardiomyocytes incubated with PMA plus various inhibitors of endocytosis-related processes: Dynasore (P+Dyn) and Pitstop (P+Pit). **(B)** Quantification of dextran internalization with PMA and the PI3K inhibitor PI103 (PI). **(C)** Isolated rat myocytes were incubated with vehicle (control), PMA or PMA+PI103 (P+PI) for 30 min, followed by the addition of PMA and further incubation for 30 min. Example blots can be seen along with a graph comparing IkB and pIkB levels. **(D)** Quantification of mean distance between t-tubules after 1 d of treatment with PMA and PI3K inhibitors wortmannin and PI103. **(E)** Quantification of dextran internalization after 2 h and of the mean distance between t-tubules after 1 d of treatment of the actin cytoskeleton inhibitor cytochalasin D (CyD) with PMA. **(F)** Quantification of dextran internalization with the FDA-approved macropinocytosis inhibitors auranofin (A) and imipramine (I) with PMA **(G)** Quantification of the mean distance between t-tubules after 1 d of treatment with myosin inhibitors: 2,3-butanedione monoxime (Bd), blebbistatin (Bl), pentabromopseudilin (Pb), and triiodophenol (Tr) with PMA.
Statistical test: paired t-test. P-values after correction for multiple comparisons: $^{*}p < 0.05$; $^{**}p < 0.01$; $^{***}p < 0.001$.

### Figure S9

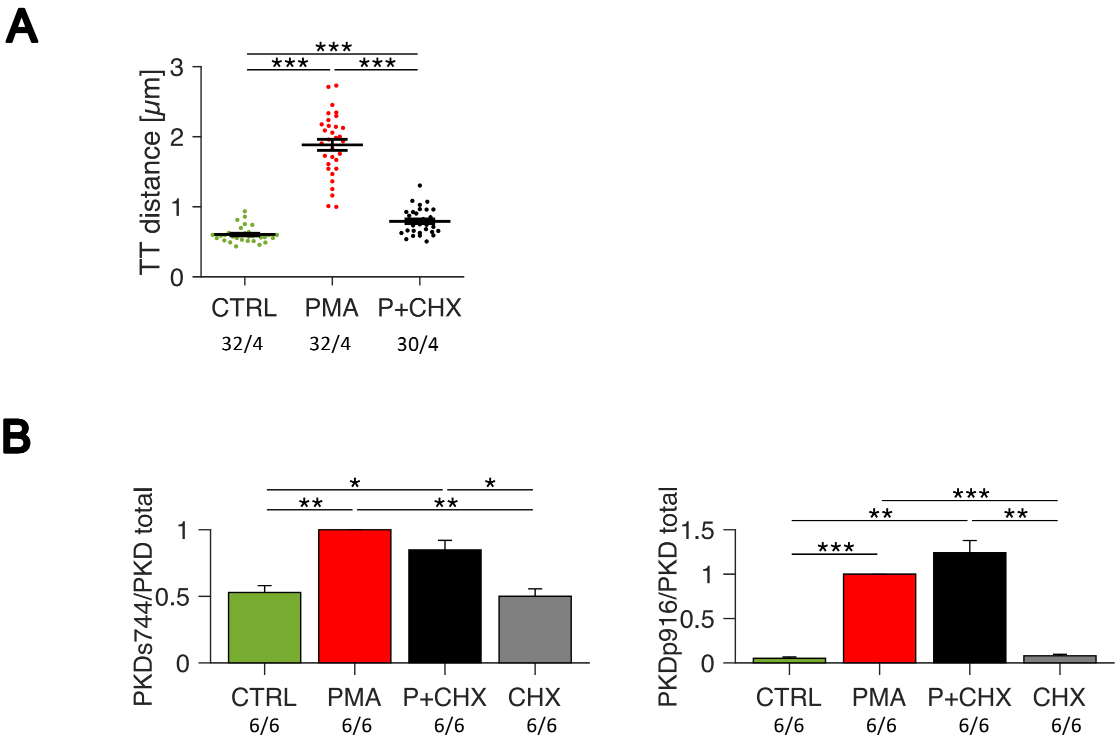


**Fig. S9. (A)** Mean t-tubular (TT) distance in rat cardiomyocytes treated with vehicle (CTRL), 50 nM PMA, or 50 nM PMA + 10 µg/mL cycloheximide (P+CHX) for 24 h. **(B)** Western Blot analysis of PKD phopshorylation at s744 and s916. * $p<0.05$, ** $p<0.01$, *** $p<0.001$; unpaired, non-equal variance (Welch's) t-test; cells from matched cell isolations.
