## Supplemental Tables for "PKC Promotes T-Tubule Membrane Loss by Activating a PKD– NFκB Endocytic Pathway"

### Supplemental Table 1

#### rat model

| Pathway | Condition | score | P value |
|---|---|---|---|
| Androgen | PMAvsCTRL | 1.127173905 | 0.259683327 |
| EGFR | PMAvsCTRL | 3.222384065 | 0.001273449 |
| Estrogen | PMAvsCTRL | -2.154014229 | 0.031251707 |
| Hypoxia | PMAvsCTRL | -1.916069586 | 0.055371366 |
| JAK-STAT | PMAvsCTRL | -4.329978469 | 1.49894E-05 |
| MAPK | PMAvsCTRL | 5.894854501 | 3.81413E-09 |
| NFkB | PMAvsCTRL | 4.344506974 | 1.40319E-05 |
| PI3K | PMAvsCTRL | 4.04653441 | 5.2189E-05 |
| TGFb | PMAvsCTRL | 3.001503628 | 0.00269002 |
| TNFa | PMAvsCTRL | -1.256570671 | 0.208924732 |
| Trail | PMAvsCTRL | -5.064264706 | 4.13851E-07 |
| VEGF | PMAvsCTRL | 3.503727161 | 0.00045986 |
| WNT | PMAvsCTRL | -0.81350309 | 0.415939964 |
| p53 | PMAvsCTRL | 3.36514573 | 0.00076656 |
| Androgen | P+TNvsPMA | -1.660820768 | 0.096766101 |
| EGFR | P+TNvsPMA | 5.695280768 | 1.25006E-08 |
| Estrogen | P+TNvsPMA | -0.629993799 | 0.528706287 |
| Hypoxia | P+TNvsPMA | 3.705198548 | 0.00021183 |
| JAK-STAT | P+TNvsPMA | -1.711352588 | 0.087032516 |
| MAPK | P+TNvsPMA | -3.550752111 | 0.000385069 |
| NFkB | P+TNvsPMA | -5.953776992 | 2.66675E-09 |
| PI3K | P+TNvsPMA | -4.743978902 | 2.11098E-06 |
| TGFb | P+TNvsPMA | 1.640412521 | 0.10093613 |
| TNFa | P+TNvsPMA | 0.202778656 | 0.839310233 |
| Trail | P+TNvsPMA | 0.222630038 | 0.823825871 |
| VEGF | P+TNvsPMA | -1.719075564 | 0.085617052 |
| WNT | P+TNvsPMA | 3.717111869 | 0.000202097 |
| p53 | P+TNvsPMA | -0.920604699 | 0.357268644 |
| Androgen | P+TNvsCTRL | 0.060763146 | 0.951548486 |
| EGFR | P+TNvsCTRL | 6.333550813 | 2.45007E-10 |
| Estrogen | P+TNvsCTRL | -2.389290107 | 0.016890749 |
| Hypoxia | P+TNvsCTRL | 0.339131465 | 0.734514479 |
| JAK-STAT | P+TNvsCTRL | -5.002472796 | 5.71091E-07 |
| MAPK | P+TNvsCTRL | 3.575907402 | 0.000349889 |
| NFkB | P+TNvsCTRL | 0.695656351 | 0.486652654 |
| PI3K | P+TNvsCTRL | 1.11039935 | 0.26684116 |
| TGFb | P+TNvsCTRL | 3.684445648 | 0.000229843 |
| TNFa | P+TNvsCTRL | -1.005656399 | 0.314593809 |
| Trail | P+TNvsCTRL | -4.707754228 | 2.52244E-06 |
| VEGF | P+TNvsCTRL | 2.243292948 | 0.024889511 |
| WNT | P+TNvsCTRL | 1.283447143 | 0.199351264 |
| p53 | P+TNvsCTRL | 2.718165116 | 0.006570516 |

#### human model

| Pathway | Condition | Score | P value |
|---|---|---|---|
| Androgen | PMAvsCTRL | 3.602796439 | 0.000315623 |
| EGFR | PMAvsCTRL | 11.1290986 | 1.11269E-28 |
| Estrogen | PMAvsCTRL | -0.994965028 | 0.319766062 |
| Hypoxia | PMAvsCTRL | -2.853632669 | 0.004326962 |
| JAK-STAT | PMAvsCTRL | -5.892410009 | 3.8709E-09 |
| MAPK | PMAvsCTRL | 10.34703173 | 5.0376E-25 |
| NFkB | PMAvsCTRL | 6.153046494 | 7.75442E-10 |
| PI3K | PMAvsCTRL | 8.58972174 | 9.38808E-18 |
| TGFb | PMAvsCTRL | 5.731634191 | 1.00989E-08 |
| TNFa | PMAvsCTRL | -1.462158175 | 0.143714562 |
| Trail | PMAvsCTRL | -6.006693015 | 1.92825E-09 |
| VEGF | PMAvsCTRL | 9.492656601 | 2.51373E-21 |
| WNT | PMAvsCTRL | -3.250338707 | 0.001154701 |
| p53 | PMAvsCTRL | 5.916360223 | 3.34844E-09 |
| Androgen | P+TNvsPMA | -2.769564432 | 0.005618626 |
| EGFR | P+TNvsPMA | 4.800344981 | 1.59608E-06 |
| Estrogen | P+TNvsPMA | -1.526122713 | 0.126996105 |
| Hypoxia | P+TNvsPMA | 1.091696529 | 0.274980482 |
| JAK-STAT | P+TNvsPMA | -9.270746001 | 2.04302E-20 |
| MAPK | P+TNvsPMA | -3.593561422 | 0.000327021 |
| NFkB | P+TNvsPMA | -9.005557456 | 2.34599E-19 |
| PI3K | P+TNvsPMA | -6.268797992 | 3.71757E-10 |
| TGFb | P+TNvsPMA | 0.542603226 | 0.58740944 |
| TNFa | P+TNvsPMA | -4.22964583 | 2.35165E-05 |
| Trail | P+TNvsPMA | 3.369598615 | 0.000754288 |
| VEGF | P+TNvsPMA | -4.46277626 | 8.13737E-06 |
| WNT | P+TNvsPMA | 3.378976062 | 0.000729039 |
| p53 | P+TNvsPMA | -0.552405361 | 0.580677216 |
| Androgen | P+TNvsCTRL | 1.795447324 | 0.07259849 |
| EGFR | P+TNvsCTRL | 13.25532298 | 6.34627E-40 |
| Estrogen | P+TNvsCTRL | -1.834483883 | 0.06659797 |
| Hypoxia | P+TNvsCTRL | -1.976997015 | 0.048056534 |
| JAK-STAT | P+TNvsCTRL | -10.51591713 | 8.60295E-26 |
| MAPK | P+TNvsCTRL | 7.659308639 | 1.95962E-14 |
| NFkB | P+TNvsCTRL | 0.722745222 | 0.469845405 |
| PI3K | P+TNvsCTRL | 4.403001677 | 1.07351E-05 |
| TGFb | P+TNvsCTRL | 5.703250541 | 1.19307E-08 |
| TNFa | P+TNvsCTRL | -3.696172177 | 0.000219495 |
| Trail | P+TNvsCTRL | -3.806094516 | 0.000141625 |
| VEGF | P+TNvsCTRL | 6.267540643 | 3.74764E-10 |
| WNT | P+TNvsCTRL | -1.199732639 | 0.230258291 |
| p53 | P+TNvsCTRL | 5.264631541 | 1.42013E-07 |

**Supplemental Table 1.** Results from PROGENy pathway analysis, using the Wald statistics value of the differential expression analysis of PMA vs CTRL, P+TN vs PMA and P+TN vs CTRL from the rat and human models.

### Supplemental Table 2

| Group | Dextran (fraction) | SE | n/N | TT distance [µm] | SE | n/N |
|---|---|---|---|---|---|---|
| 2,3-Butanedione (BDM) | 0.0014 | 0.000265 | 30/4 | 0.43 | 0.013 | 14/3 |
| BDM+PMA | 0.0036 | 0.000540 | 30/4 | 0.54 | 0.017 | 40/9 |
| Angiotensin+Endothelin+Phenylephrine | 0.0063 | 0.000246 | 81/8 | 0.62 | 0.016 | 51/6 |
| Blebbistatin+PMA | 0.0070 | 0.000552 | 31/4 | 0.98 | 0.080 | 32/6 |
| BMS+PMA | 0.0031 | 0.000896 | 26/3 | 0.90 | 0.061 | 28/4 |
| Control (CTRL) | 0.0042 | 0.000166 | 264/34 | 0.64 | 0.007 | 802/91 |
| CRT0066101+PMA | 0.0061 | 0.000577 | 32/4 | 0.54 | 0.025 | 38/5 |
| Cycloheximide (CHX)+PMA | 0.0045 | 0.000479 | 24/3 | 0.70 | 0.039 | 16/2 |
| Cytochalasin D+PMA | 0.0082 | 0.000343 | 60/6 | 1.04 | 0.088 | 32/4 |
| GÖ6983+PMA | 0.0045 | 0.000707 | 8/1 | 0.70 | 0.045 | 26/4 |
| Imipranine+PMA | 0.0088 | 0.000328 | 60/6 | 1.64 | 0.089 | 31/4 |
| KU-55933+PMA | 0.0078 | 0.000461 | 24/3 | 1.00 | 0.132 | 32/4 |
| PI103+PMA | 0.0055 | 0.000453 | 39/5 | 0.61 | 0.029 | 39/5 |
| PMA | 0.0085 | 0.000240 | 264/34 | 1.44 | 0.019 | 727/91 |
| TPCA1+NIKSMI+PMA | 0.0068 | 0.000556 | 58/7 | 0.59 | 0.027 | 44/6 |
| Wortmannin+PMA | 0.0062 | 0.000651 | 24/3 | 0.89 | 0.079 | 32/4 |

**Supplemental Table 2: Correlation of internalization of dextran and t-tubule distance**

The table displays the mean, standard error and n/N (cells/animals) for the values plotted in Figure 7F. For the endocytosis assay, isolated rat cardiomyocytes were incubated with vehicle (CTRL) or the inhibitor for 1 h, followed by the addition of 50 nM PMA and further incubation for 2 h. The combination of angiotensin, endothelin and phenylephrine was incubated for a total of 2 h. Blebbistatin, a fast-acting myosin II inhibitor, was added 10 min before the end of the incubation (after 1 h with vehicle and 2 h with PMA). For the t-tubule assay (TT distance), rat myocytes were kept in culture for a total time of of one day. Vehicle (control) or an inhibitor were added to the culture medium immediately after cell isolation. One hour later, PMA was added. After approximately 24 h, the membrane was stained with Di8-ANEPPS.

### Supplemental Table 3

| Target | 1st AB | 1st AB Host | 1st AB solution | 2nd Ab | 2nd AB Host | 2nd AB solution |
|---|---|---|---|---|---|---|
| PKD total | Abcam, ab51246 | Rabbit | 1:1000, 1% Milk, 0.05% Tween, in TBS-buffer, 12H, 4°C | a-Rabbit HRP (Invitrogen, G21234) | Goat | 1:40000, 1% Milk / 2.5% BSA, 1H, RT |
| PKD p-S916 | Cell Signalling, 2051 | Rabbit | 1:1000, 1x Rotiblock, 0.05% Tween, in TBS-buffer, 12H, 4°C | a-Rabbit HRP (Invitrogen, G21234) | Goat | 1:40000, 1% Milk / 2.5% BSA, 1H, RT |
| PKD p-S744-748 | Cell Signalling, 2054 | Rabbit | 1:1000, 1% Milk, 0.05% Tween, in TBS-buffer,12H, 4°C | a-Rabbit HRP (Invitrogen, G21234) | Goat | 1:40000, 1%Milk/2.5%BSA, 1H, RT |
| IkBa | Cell Signalling, 9242 | Rabbit | 1:1000, 1% Milk, 0.05% Tween, in TBS-buffer, 12H, 4°C | a-Rabbit HRP (Invitrogen, G21234) | Goat | 1:40000, 1%Milk/2.5%BSA, 1H, RT |
| p-IkBa | Cell Signalling, 9246 | Mouse | 1:1000, 5% Milk, 0.05% Tween, in TBS-buffer, 12H, 4°C | a-Mouse HRP (Abcam, ab97023) | Goat | 1:50000, 5% Milk, 1H, RT |

**Supplemental Table 3: Western Blot Antibodies**

### Supplemental Table 4

| Target-Fluorophore | Fixation | Blocking Solution | 1st AB | 2nd AB |
|---|---|---|---|---|
| • DAPI<br>• LTCC-AF555<br>• RyR-AF488<br>• JPH2-AF647 | PFA 2%<br>RT<br>5 min | 5% BSA<br>5% NGS<br>0.25% Triton-X in PBS | • a-CACNA1C, 1:200, Guinea Pig polyclonal (Alomone Labs, ACC-003-GP)<br>• a-RyR, 1:200, Mouse IgG1 (Thermo Fisher, MA3916)<br>• a-JPH2, 1:200, Rabbit polyclonal (Thermo Fisher, 40-5300) | • DAPI ()<br>• a-MouseIgG1-AF488, 1:200, Goat (Invitrogen, A21121)<br>• a-GuineaPigH+L-AF555, 1:200, Goat (Abcam, ab150186)<br>• a-RabbitH+L-AF647, 1:200, Goat (Invitrogen, A32733) |
| • DAPI<br>• BIN1-AF488 | Acetone<br>4°C<br>7 min | PBS | • a-Amphiphysin II, 1:100, Mouse IgG1 (Santa Cruz, sc-23918) | • DAPI ()<br>• a-MouseIgG1-AF488, 1:200, Goat (Invitrogen, A21121) |

**Supplemental Table 4: Antibodies / Dyes for Fluorescent Imaging**

### Supplemental Table 5

| Staining | Fluorophore | Excitation (nm) | Detection (nm) |
|---|---|---|---|
| **Membrane** | Di-8-ANEPPS | 488 | 500 - 758 |
| | FM 4-64 | 561 | 592 - 759 |
| | Hoechst | 405 | 373 - 483 |
| | Lipofuscin | 488 | 580 - 620 |
| **Dextran assay** | AF 488 | 488 | 489 - 577 |
| | AM 4-64 | 561 | 650 - 751 |
| | Lipofuscin | 633 | 700 - 800 |
| **DAPI / LTCC / RyR / JPH2** | DAPI | 405 | 418 - 554 |
| | AF488 | 488 | 495 - 556 |
| | AF555 | 561 | 559 - 629 |
| | AF647 | 633 | 638 - 690 |
| **DAPI/BIN1** | DAPI | 405 | 423 - 476 |
| | AF 488 | 488 | 498 - 577 |

**Supplemental Table 5: Confocal Imaging Settings**
